# A gut-selective Axl Inhibitor Attenuates Intestinal Fibrosis in myofibroblast cell models and the Mouse S. Typhimurium Model *in vivo*

**DOI:** 10.1101/2025.03.30.646118

**Authors:** Yongjia Feng, Laura A. Johnson, Martin Clasby, Peter D.R. Higgins

## Abstract

**Background:** Intestinal fibrosis leads to intestinal failure in Crohn’s disease (CD), but we have no effective antifibrotic medical therapies. Axl inhibitors have been studied in the treatment of fibrosis in liver and lung, but not in intestine. The objective of this study was to test whether CCG264341, a novel gut selective Axl inhibitor, can treat intestinal fibrosis in both *in vitro* and *in vivo* IBD fibrosis models.

**Methods:** CCD-18Co human intestinal myofibroblasts were co-treated with 100 ng/mL Fas ligand (FASL) and a dose range of CCG264341 for 5 hours, or treated with TGFβ for 48 hours, followed by protein isolation and western blot analysis. Tissue PK experiments in mice were performed to select the CCG264341 dose for the *in vivo* model. For the *in vivo* model, intestinal fibrosis was induced in CBA/J mice by *S. typhimurium* infection, Mice were treated with CCG264341 daily during days 4-21 after *S. typhimurium* infection, followed by sac on day 22. Cecum and proximal colon (affected tissue) was analyzed by gross pathology, histological fibrosis scoring, qRT-PCR, western blot, and immunofluorescence staining.

**Results:** CCG264341 at doses of 0.1/0.3/1.0 µM sensitizes CCD-18Co myofibroblasts to FasL-mediated apoptosis over FasL alone, and this effect was dose dependent, per cleaved Parp protein quantitation. With TGFβ treatment, Axl protein expression was increased, along with Col1a1, MYLK, αSMA and FN1 protein upregulation. CCG264341 could reduce AXL protein and downstream signal p-Akt expression with dose dependence, along with reduced Col1a1, MYLK, αSMA and FN1 protein expression after treatment with CCG264341 treatment. The highest CCG264341 concentrations were detected in the terminal ileum after 5 hours oral feeding in PK experiments, and much lower compound concentrations could still be detected in other organs. A dose of 25mg/kg daily was selected for the mouse intestinal fibrosis treatment model. In the murine *S. typhimurium* model, AXL protein and downstream signals p-Akt, p-Erk and P-Stat3 expression were all increased in colon and cecum, with the expression of the fibrosis related proteins upregulated. CCG264341 treatment significantly inhibited AXL expression, with downstream signals and fibrosis related protein downregulation. By blinded histopathology with H&E and trichrome staining, as well as immunofluorescence for fibrosis related proteins, colon fibrosis scores significantly decreased.

**Conclusions:** CCG264341is a partially gut selective Axl inhibitor. This compound could play a therapeutic role in IBD fibrosis models. Further work to improve gut selectivity and anti-fibrotic efficacy are needed, but this is a promising pathway for future therapeutic intervention in Crohn’s disease.

## Introduction

Crohn’s disease (CD) is a chronic, progressive, and frequently debilitating intestinal disorder affecting more than 600,000 people in the US and millions worldwide. Despite the use of highly effective anti-inflammatory therapeutics, two-thirds of patients will require surgical resection of intestine in the first 30 years after diagnosis, most commonly due to fibrostenotic obstructive disease. Current mechanical treatments for fibrostenotic disease, specifically endoscopic balloon dilation, surgical strictureplasty, or surgical resection, are invasive, costly, and carry significant risks. The lack of effective anti-fibrotic drugs to treat intestinal fibrosis remains a critical treatment gap as a majority of patients progress to fibrostenotic disease and intestinal failure.

In other organ systems, the advent of antifibrotic therapeutics has overthrown the prior paradigm of the irreversibility and inexorable progression of fibrosis. Over the last decade, preclinical animal studies and human trials have identified multiple druggable targets in organ fibrosis. We have previously identified the Axl pathway as a potential target for anti-fibrotic intervention in intestinal fibrosis^1^. The Axl gene is upregulated in CD strictures, in *in vitro* models of fibrosis, and in two rodent models of intestinal fibrosis. As we have previously demonstrated, blocking AXL signaling via a systemic small molecule inhibitor (BGB324, bemcentinib) abrogates fibrosis in 3 *in vitro* models, including a complex multicellular organoid human intestinal organoid model. While the side effects of bemcentinib and other Axl inhibitors (neutropenia in 86%, diarrhea in 57%, fatigue in 57%, and nausea in 52%)^2^ are tolerable for short-term, curative chemotherapy, they are not realistic for a long-term treatment and prevention of recurrence of obstructive intestinal fibrosis in Crohn’s disease.

Axl is a receptor tyrosine kinase belonging to the TAM receptor family along with Tyro3 and MER^3,4^, and is expressed on the cell surface of mesenchymal, epithelial, and hematopoietic lineage cells^5,6^. Tissue fibrosis is characterized by extensive accumulation of extracellular matrix (ECM) and stromal elements in response to chronic inflammation and tissue injury. The TAM receptor family plays critical roles during tissue remodeling in fibrotic diseases^7,8^.Extensive studies of Axl and fibrosis have been done on several organs, especially in liver^9–11^ and lung^12–14^. In liver cirrhosis, AXL signaling is activated to induce the transcription of SLUG, a critical regulator of EMT^15^. Idiopathic pulmonary fibrosis (IPF) is another potentially deadly fibrotic disease leading to progressive loss of lung function^16^. IPF tissues express elevated levels of TAM receptors and GAS6, and small-molecule inhibitors that target multiple TAM receptor kinases exhibit enhanced anti-fibrotic activity^17^, suggesting that IPF has activated the entire TAM receptor family. Based on the demonstrated benefit of Axl inhibitors on liver and lung fibrosis, Axl is a promising target for the treatment of intestinal fibrosis in Crohn’s disease.

Fibrosis in Crohn’s disease is characterized by an excessive wound healing response, initiated and propagated by apoptosis resistant myofibroblasts^18–20^. Few studies exist for the role of AXL in IBD related fibrosis. Our published and current data demonstrate that TGFβ significantly increases the expression of Axl in CCD18-Co myofibroblasts as well as in human intestinal organoids, and BGB324, a commercial Axl inhibitor, could block TGFβ-induced fibrosis and increased the levels of cleaved-PARP protein in these cells corresponding to induction of myofibroblast apoptosis^1^. However, BGB324 failed to prevent the cecal and proximal colon fibrosis in *Salmonella typhimurium* infected mice, which may be due to the low concentration in gut and toxicity at higher doses. This led us to develop an Axl inhibitor with strong gut selectivity, to enhance drug efficacy and reduce toxicity.

Gut-selective drugs provide excellent cell permeability for rapid gut uptake after oral administration but produce low plasma levels after first-pass metabolism in the liver. This strategy has been used for the development of gut-selective prodrugs like budesonide^21^. Based on our previous data, we worked to develop a novel gut-selective Axl inhibitor for anti-fibrotic treatment in patients with Crohn’s disease.

## Materials and Methods

### TGF-β1 Fibrogenesis Cell Model

Human colonic myofibroblast CCD-18co cells (CRL-1459) were obtained from ATCC (Bethesda, MD). Low-passage (passages 3–10) colonic myofibroblasts (CCD-18co cells) were seeded at 5 × 10^4^ cells/mL on multi-well tissue culture plates (12-well, 1 mL/well for gene expression; 6-well, 3 mL/well for protein expression) in α-MEM medium (Invitrogen, Carlsbad, CA). Cells were serum-starved overnight before stimulation with 0.05 ng/mL of TGF-β1 (R&D Systems, Minneapolis, MN), and cotreated +/- CCG264341 at a range of concentrations (0/0.1/0.3/1.0µM). The maximum concentration of DMSO in any *in vitro* experiment was ≤0.1%. Serum starvation was performed to reduce the effect of serum on markers of fibroblast differentiation during treatment with TGF-β1. Cells were harvested for gene expression analysis at 24 hours and for protein expression analysis at 48 hours^1,22,23^.

### Fas Ligand–mediated Apoptosis Assay

Low-passage (passages 2-10) CCD-18co cells were seeded at 1 × 10^5^ cells/mL in 10% FBS-containing α-MEM growth media (Invitrogen, Carlsbad, CA) on multi-well plates, Media was aspirated off the plastic wells and replaced with low-serum (0.5% FBS) α-MEM for overnight starvation. The following day, apoptosis was induced by treatment with 100 ng/mL Fas activating antibody (clone CH11, Millipore Sigma, St. Louis, MO) (FASL) in the presence or absence of a range of concentrations of CCG264341 (0.1/0.3/1.0µM). At 5 hours, the total cells were washed, scraped, and collected, cell were homogenized and sonicated with RAPI buffer and cell lysates were used for apoptosis-related protein cleaved-Parp analysis^1,23,23^.

### SiRNA transfection

AXL Human siRNA Oligo Duplex (Locus ID 558) were purchased from Origene company (SKU SR319445), the transfection was processed according to the manufacturer’s protocol^24^, Briefly, in a six well tissue culture plate, seed 5 × 10^4^ cells/mL CCD-18co cells in 3 ml antibiotic-free normal growth medium supplemented with FBS, incubate the cells at 37° C in a CO2 incubator for 24 hours until the cells are 60-80% confluent. To prepare transfection solution, make 2 1.5ml EP tubes, solution A: For each transfection, dilute 50pmols siRNA into 100 µl siRNA Transfection Medium; Solution B: For each transfection, dilute 4 µl of siRNA Transfection into 100 µl siRNA Transfection Medium. Add the siRNA duplex solution (Solution A) directly to the dilute Transfection Reagent (Solution B), mix gently by pipetting the solution up and down and incubate the mixture 15-45 minutes at room temperature. For each transfection, add 0.8 ml siRNA Transfection Medium to each tube containing the siRNA Transfection Reagent mixture (Solution A + Solution B). Mix gently and overlay the mixture onto the cells, incubate the cells 5-7 hours at 37° C in a CO2 incubator. Add 1 ml of normal growth medium containing 2 times the normal serum concentration (2x normal growth medium) without removing the transfection mixture, incubate the cells for an additional 24 hours, replace with fresh FBS free medium and start TGF-β1 or FasL stimulation as prescribed.

### Mice

Female 8 to 10-week-old CBA/J mice (Jackson Laboratories, Bar Harbor, ME) received 20 mg streptomycin on day 0 by oral gavage to eradicate the commensal microbiota 24 hours prior to *S. typhimurium* infection. Stool cultures were used to make sure the commensal microbiota was eradicated. Each mouse received 3×10^6^ cfu *S. typhimurium* strain SL1344 in 100µl 0.1 M HBSS buffer (pH = 8.0) by oral gavage on day 1. Control mice received same amount of 0.1 M HBSS. Stool culture was performed to make sure the *S. typhimurium* infection was successful^25,26^. Four days after *S. typhimurium* infection, mice were dosed with 25 mg/kg/day CCG264341 (100µL) by oral gavage daily until day 21. At the same time, the control group was given the vehicle (ETOH: PEG400: Phosal50 10:30:60). The mice were divided to 4 treatment groups: group 1: *S. typhimurium* (-) CCG264341(-)*; group 2: S. typhimurium* (-) CCG264341(+)*; group 3: S. typhimurium* (+) CCG264341(-)*; group 4 S. typhimurium* (+) CCG264341(+) (Figure 7A), and humane euthanization was performed at day 22 post-infection before the analysis of intestinal tissues to determine the effect of CCG264341 on intestinal fibrosis.

### Gross pathology and tissue collection

After mice were euthanized, the caecum and proximal colon were collected, photographed, measured, and weighed. The cecal two-dimensional area was determined from digital photographs using ImageJ ROI to delineate the cecum. To control for differences in focal distance, cecal area was normalized against a 1 cm marker in the photographic image. Tissues were snap-frozen in liquid nitrogen and stored at -80°C before molecular analysis. Cecal contents were collected and serially diluted before plating onto LB/streptomycin plates to determine *S. typhimurium* bacterial titers.

### Stool Cultures

A fresh stool was collected, suspended in 500 µL sterile PBS, streaked onto LB plates containing 10 µg/ml streptomycin, and cultured overnight at 37°C to make sure the infection was successful.

### Histology and Fibrosis score Assessment

Formalin-fixed and paraffin-embedded tissues (cecum and proximal colon) were stained with hematoxylin and eosin (H&E) and Masson’s trichrome by the University of Michigan Rogel Cancer Center Tissue and Molecular Pathology (TMP) Core [Ann Arbor, MI]. Digital photomicrographs of tissue sections were taken with an Olympus BX microscope [University of Michigan Microscopy and Image Analysis Laboratory]. A blinded pathologist performed histological fibrosis scoring. Inflammation and epithelial damage were determined using a Wirtz Scale scoring system (Table 1) in H&E-stained slides, Fibrosis was scored (0-4) according to the collagen deposition in the Masson’s trichrome stained slides Tissue measurements were quantitated with ImageJ analysis software [NIH, Bethesda, MD].

### Western immunoblotting

The collected cells or mouse cecal tissues were lysed in Pierce IP lysis buffer (ThermoFisher). The insoluble materials in the lysates were removed by centrifugation at 14,000 × g for 10 min at 4°C. Protein quantification was performed using a standard Pierce BCA protein Assay kit (Fisher Science, Rockford, IL). The lysate proteins were resolved by 4–20% sodium dodecyl sulfate polyacrylamide gel electrophoresis followed by polyvinylidene difluoride membrane transfer. The transferred membrane was probed with primary antibodies (Table 1), followed by the addition of the corresponding horseradish peroxidase-conjugated secondary antibodies. Protein bands were visualized by using ECL reagents and exposed to radiographic film. Bands were quantified using ImageJ and normalized to GAPDH and the protein/GAPDH ratio was calculated for each lane.

### Immunofluorescent staining

8 well Lab-tek II chamber slides were used for CCD-18co cell culture, and 0.5 × 10^4^ cells in 500 µL of media per well were seeded, and treatment was as described in the TGFβ or FASL models described below. At the endpoint of experiments, the cells were fixed with cold 4% PFA for 15 minutes, and washed 3 times with PBS, 10 minutes for each wash, then the chambers were stored in -20 degree for future staining.

The mouse cecum was collected in 4% PFA overnight at 4°C, then the tissues were transferred to 20% sucrose in 4°C until all of the tissues were seeded to the tube bottom. Then the tissues were embedded in OCT, and the OCT blocks were cut to 5 µmeter section width using a Leica CM1900 cryostat.

For the staining, the chamber cells or tissue slides were incubated with primary antibodies overnight in 4 °C, washed with PBS 3 times x 3 minutes, then incubated with the corresponding secondary antibodies for 1 hour at room temperature in the dark. After 3 more e minute washes with PBS, the slides were mounted with Prolong gold antifade reagent with DAPI (Invitrogen). These slides were observed, and picture were taken using Nikon Eclipse Te2000-s microscope.

### Quantitative real-time polymerase chain reaction

RNA was extracted from the cells using the RNeasy kit (Qiagen, Valencia, CA) or from the colon and cecum using the Trizol reagent (Invitrogen). Reverse transcription of 2 µg of total RNA was performed with the Superscript First Strand RT kit [Invitrogen]. Quantitative real-time polymerase chain reaction (qRT-PCR) was performed using a Stratagene Mx3000P real-time PCR system (Stratagene, La Jolla, CA). Cycling conditions were 95°C for 10 min, followed by 40 cycles of 95°C for 15 s and 62°C for 60 s. Gene expression was normalized to GAPDH as the endogenous control and fold-changes relative to uninfected controls were calculated using the ΔΔCt-method.

### Statistical Analysis

Statistical differences were determined by a 2-sided, unpaired Student *t* test or Mann-Whitney *U* test, where appropriate. Results with a *P* value <0.05 were considered statistically significant.

## Results

### Screening of many candidate compounds for in vitro inhibition of target protein expression led to the selection of CCG264341(C341) as the lead compound

Based on the backbone of BGB324, 10 variations of this were developed aimed at reducing systemic exposure, and the anti-fibrotic efficacy were tested *in vitro* in our TGF-β1 CCD-18co fibrogenesis model across a range of concentrations. Results of real-time PCR showed that among these 10 compounds, CCG254341, CCG265467, CCG365347 and 270841 showed the strongest anti-fibrotic efficacy by fibrosis-related gene expression (Figures 1 and 2), and further experiments were performed to confirm the anti-fibrotic efficacy on protein expression with western blotting (Figure 3). The protein data showed that compared to BGB324, CCG264341 showed the strongest AXL inhibition by reducing p-Akt expression and by decreasing the expression of fibrosis-related proteins (col1a1, αSma, Fn1).

**Figure 1.**
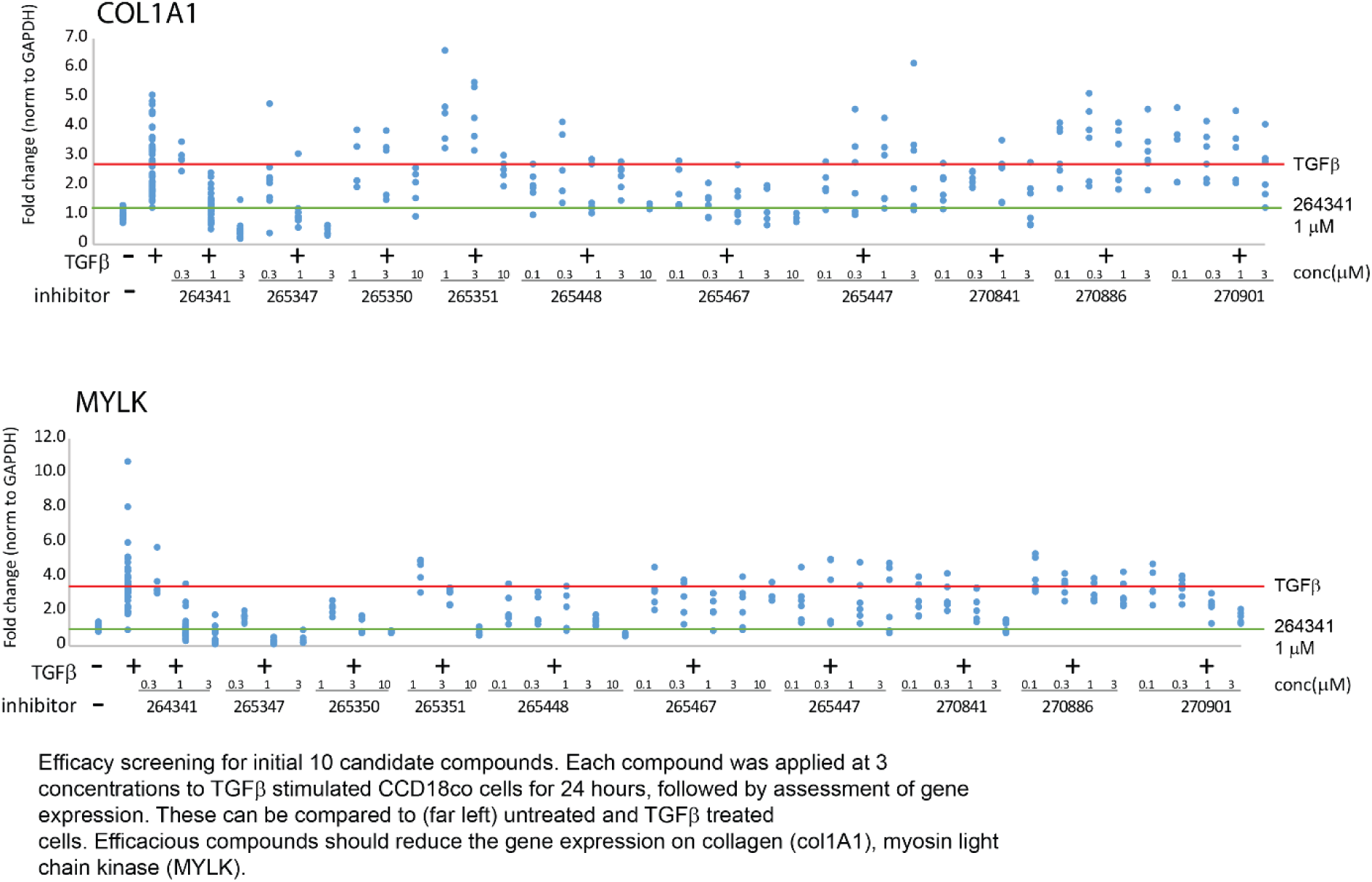
Efficacy screening for initial 10 candidate compounds. Each compound was applied at 3 concentrations to TGFβ stimulated CCD18co cells for 24 hours, followed by assessment of gene expression. These results can be compared to (far left) untreated and TGFβ treated cells. Efficacious compounds should reduce the gene expression of collagen 1 (Col1a1) and myosin light chain kinase (MYLK)

**Figure 2.**
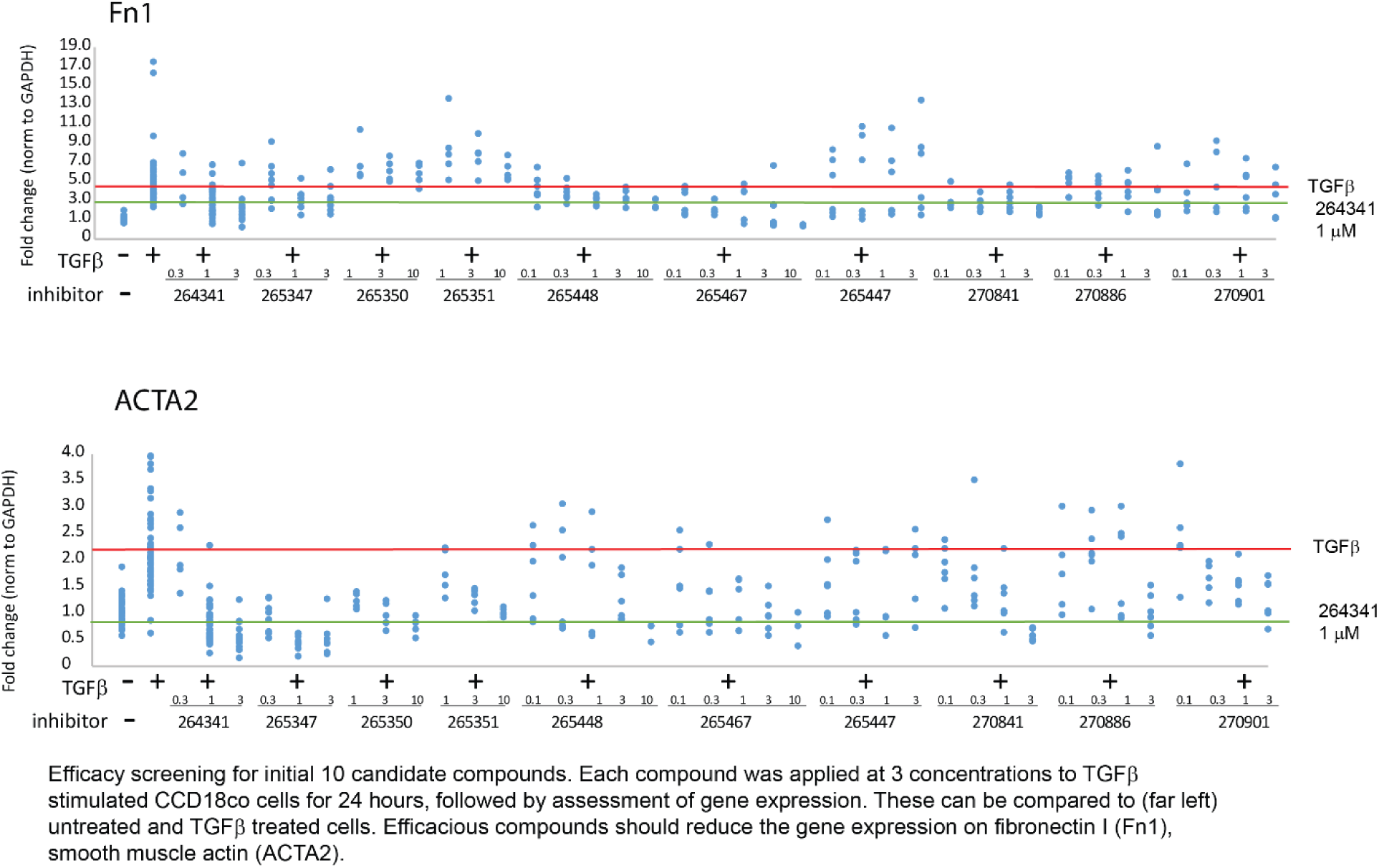
Efficacy screening for initial 10 candidate compounds. Each compound was applied at 3 concentrations to TGFβ stimulated CCD18co cells for 24 hours, followed by assessment of gene expression. These results can be compared to (far left) untreated and TGFβ treated cells. Efficacious compounds should reduce the gene expression of fibronectin and alpha smooth muscle actin (ACTA2)

**Figure 3.**
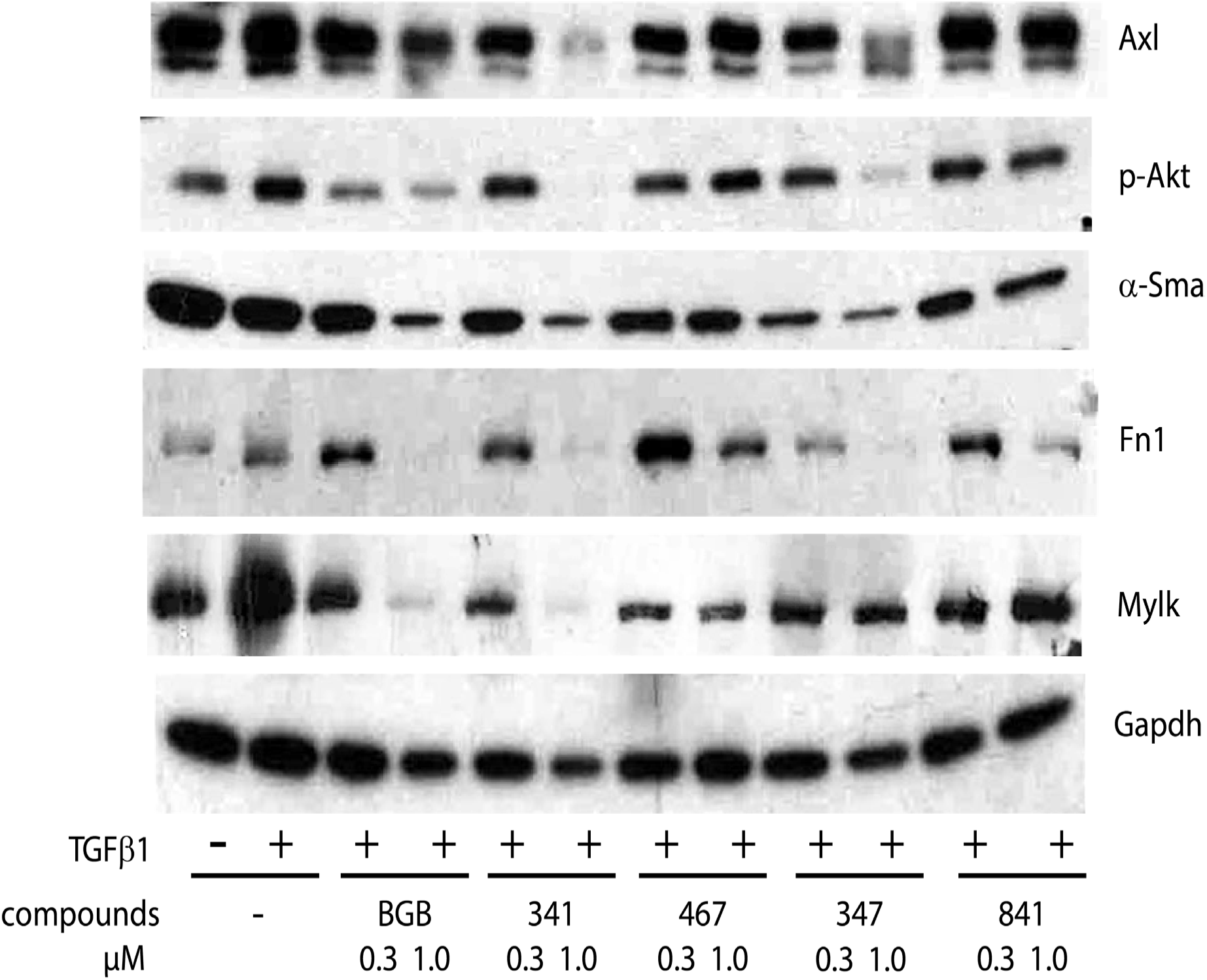
Western blot of CCD18co cell lysates after treatment with TGFβ (fibrogenic) and low vs. high doses of promising candidate compounds for 48 hours.

The metabolic stability of our candidate compounds was evaluated with mouse and human liver microsomes supplemented with NADPH (Figure 4). These results showed that CCG264341 and CCG265467 have less microsomal stability than the other candidates, which is beneficial for rapid first-pass metabolism in the liver, which increases gut selectivity. The PK experiment in mice (Figure 5) showed that high concentrations of CCG264341 were detected in gut tissue, and the peak concentration was at 5 hours after treatment. CCG264341 concentrations were measured 5 hours after treatment at 30 and 100mg/kg dose (Figure 6), high dose did show higher concentration, but not 3 folds high. Based on above data, CCG264341 were selected as the lead compound for further evaluation.

**Figure 4.**
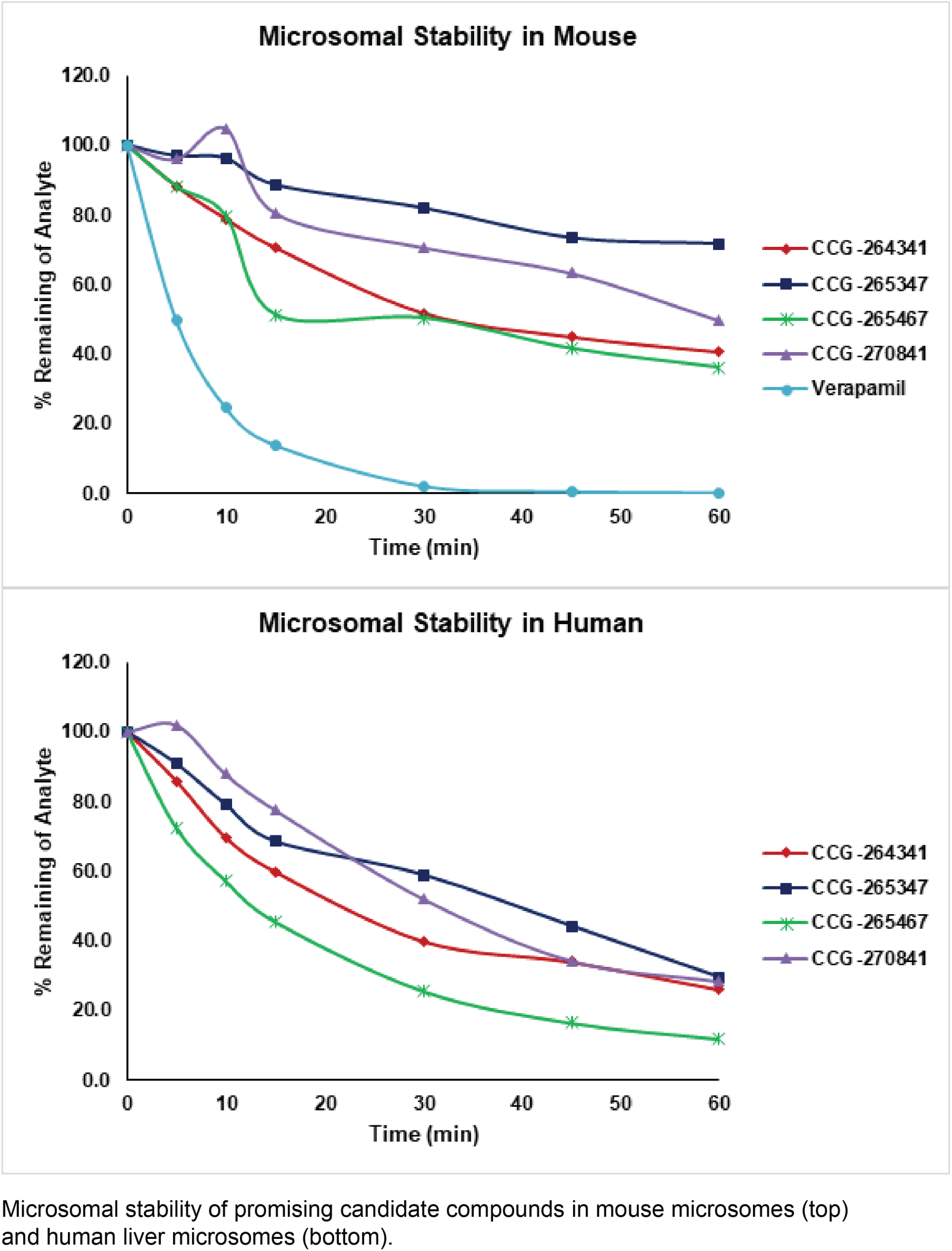
Testing microsomal stability of promising candidate compounds in mouse (upper panel) and human (lower panel). Rapid degradation by the liver will enhance candidate compound breakdown and gut selectivity.

**Figure 5.**
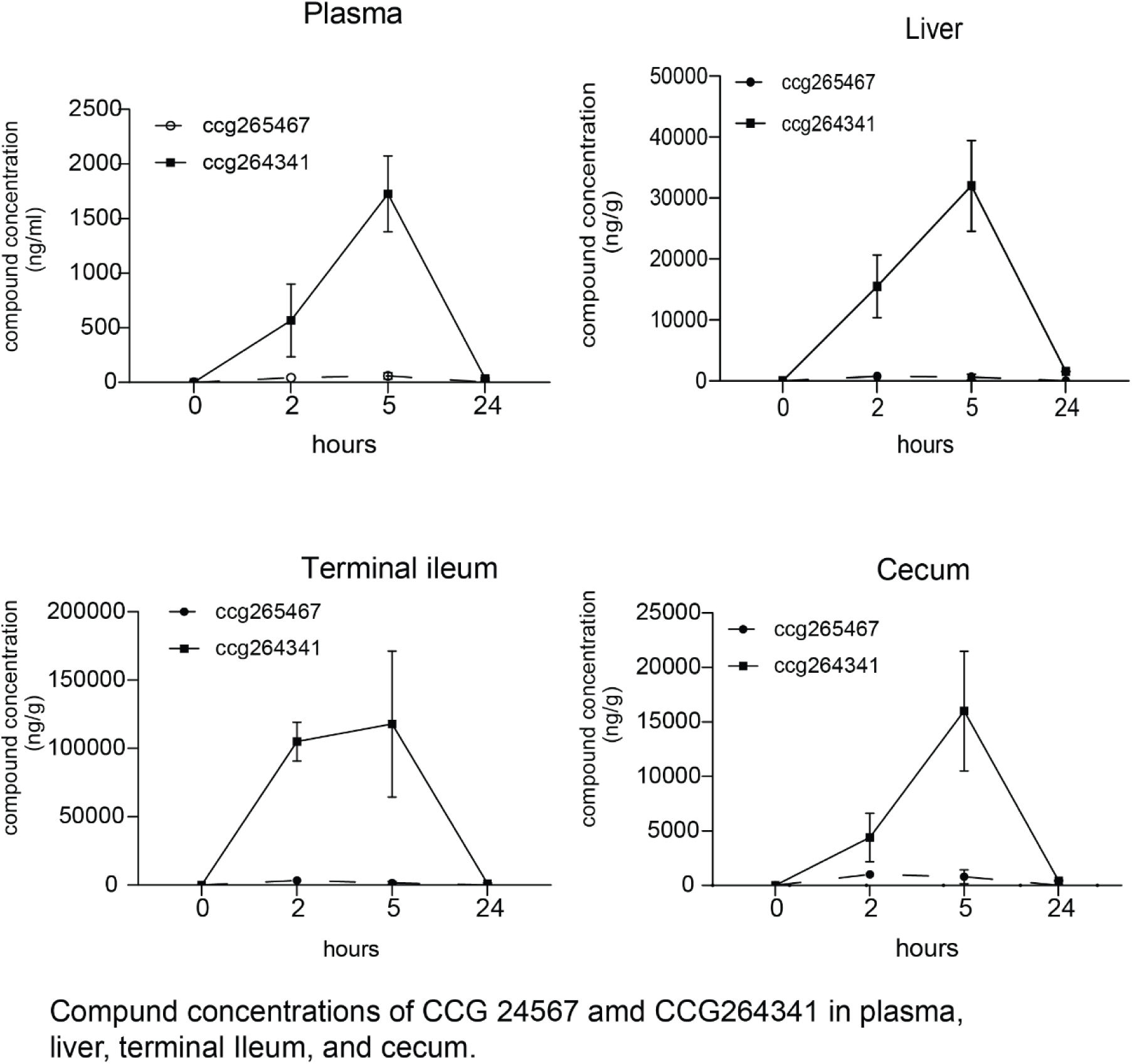
Testing tissue concentrations of promising candidate compounds in mouse plasma, liver, terminal ileum, and cecum. CCG24341 is more gut-selective than CCG265467.

**Figure 6.**
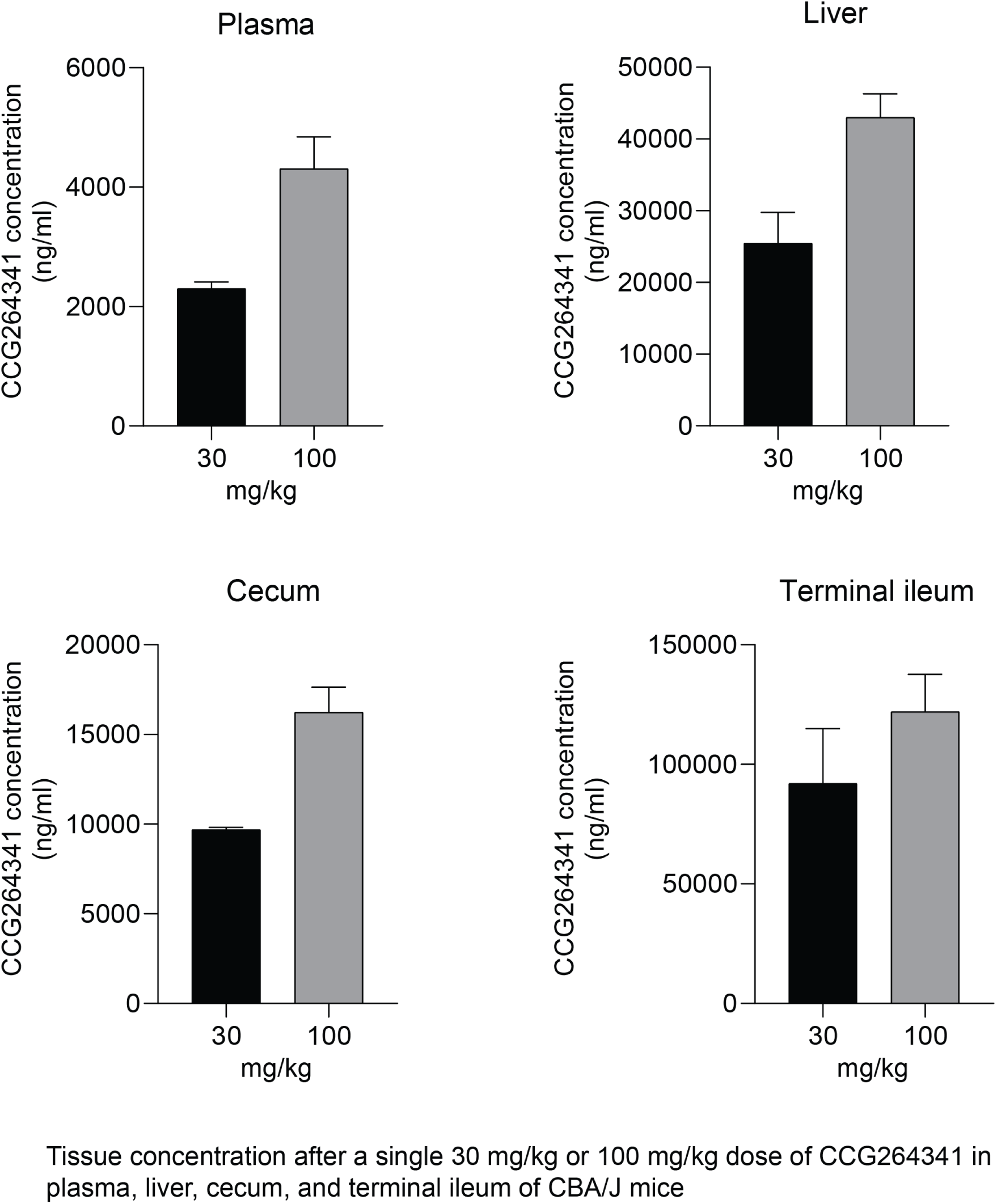
Testing tissue concentrations after a single 30 mg/kg or 100 mg/kg dose of CCG264341 in plasma, liver, cecum, and terminal ileum of CBA/J mice.

### CCG264341 Reduced Fibrosis in the *in vitro* TGF-β–mediated fibrogenesis model, and the efficacy was comparable to Axl siRNA knockdown

To confirm the anti-fibrotic efficacy with Axl inhibition, AXL protein expression was knocked down with siRNA (Figures 7 and 8). These results showed that compare to lipofectin controls and scrambles siRNA controls, AXL protein expression was successfully knocked down by AXL specific siRNA transfection, with reduction of downstream p-Akt expression, and downregulation of fibrosis related protein (Col1a1, aSMA, MYLK and FN1) expression. Immunofluorescence staining (Figure 9) data illustrates these results, further demonstrating that AXL inhibition could play a significant role in treating intestinal fibrosis.

**Figure 7.**
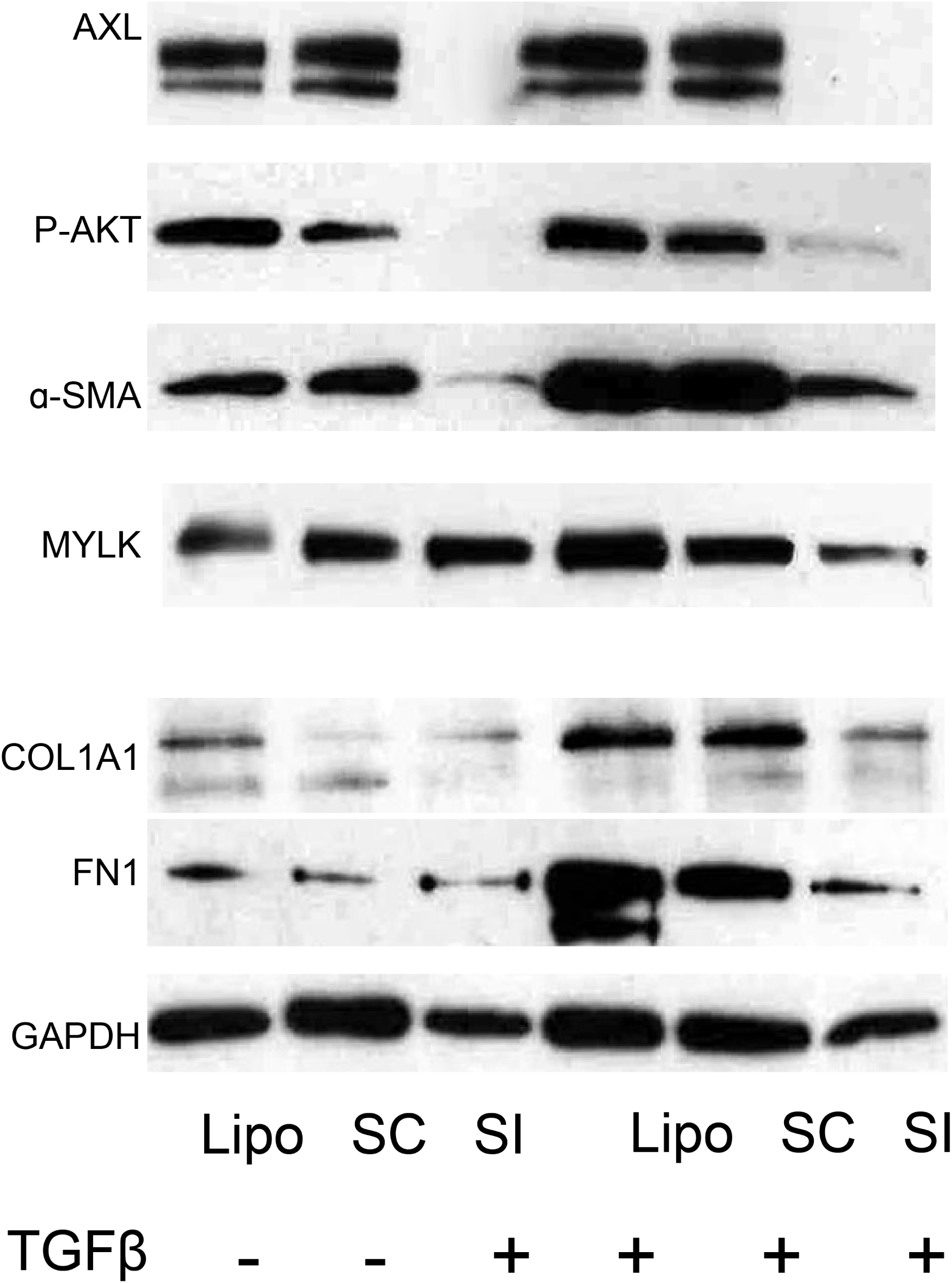
Western blotting of fibrogenic proteins, including AXL, pAkt, aSMA, MYLK, collagen 1, and fibronectin in lysates from CCD18 co human myofibroblasts treated with lipofectamine, scrambled RNA, or siRNA, and in the absence/presence of TGFβ to induce fibrogenesis.

**Figure 8.**
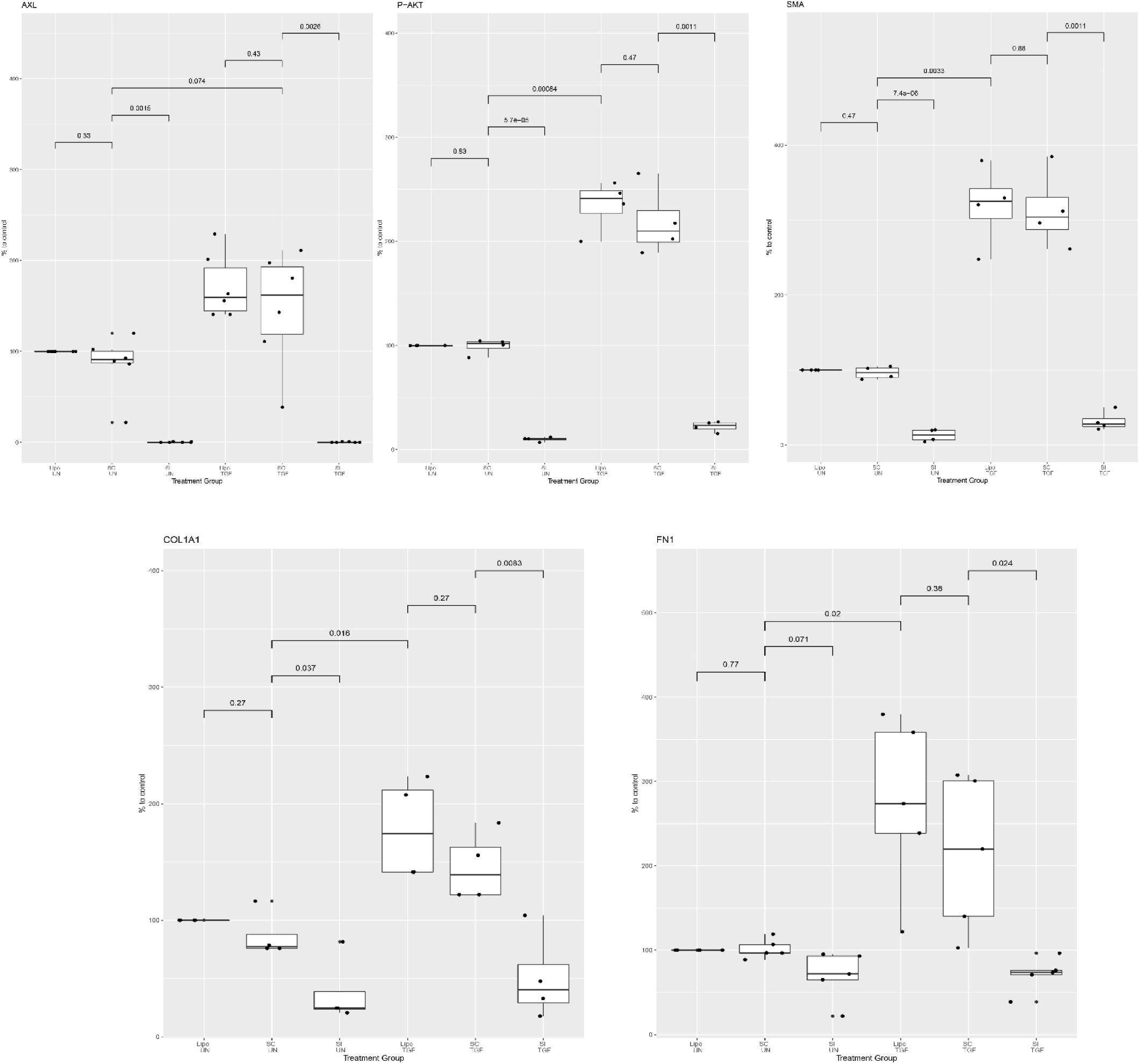
Plots of Gene expression of Axl, pAkt, alpha SMA, Collagen 1, and Fibronectin in CCD18co cells treated with siRNA vs lipofectin vs scrambled RNA in the presence or absence of TGFβ. The y axis is gene expression as a percentage of the control.

**Figure 9.**
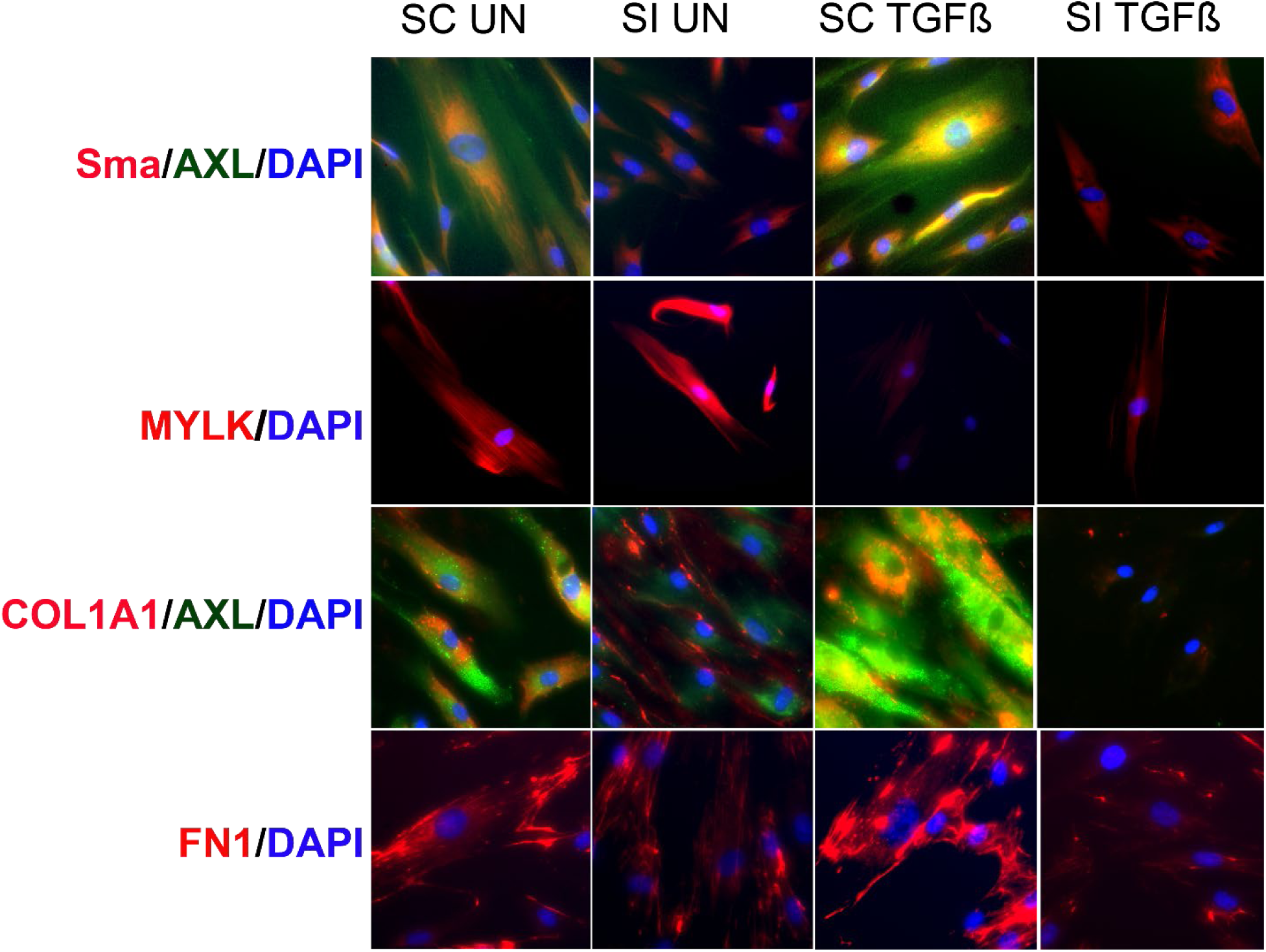
Immunofluorescence of CCD18co cells treated with AXL siRNA (vs. scrambled) and TGFβ. Antibodies to Ama, AXL, DAPI, MYLK, collagen, and fibronectin are shown. Knockdown of AXL dramatically reduces the expression of fibrogenic proteins, even in the presence of TGFβ.

We then tested the anti-fibrotic efficacy of the lead compound CCG264341 in the CCD18co TGFβ fibrogenesis cell model. Our results showed that TGFβ treatment increases AXL and downstream signal p-AKT expression, while CCG264341 treatment partially blocked both AXL and p-Akt expression in a dose-dependent manner (Figure 10). At the highest dose used, 1.0 μm of CCG264341 reduced AXL and p-Akt expression levels to baseline (Figure 11). Fibrosis related protein levels were measured (Figures 12 & 13), and we found that TGFβ stimulation increased fibrosis related proteins (Col1a1, αSMA, MYLK, and FN1) expression, and CCG264341 reduced all of these proteins in a dose-dependent manner. Immunofluorescence staining confirmed these results (Figure 14).

**Figure 10.**
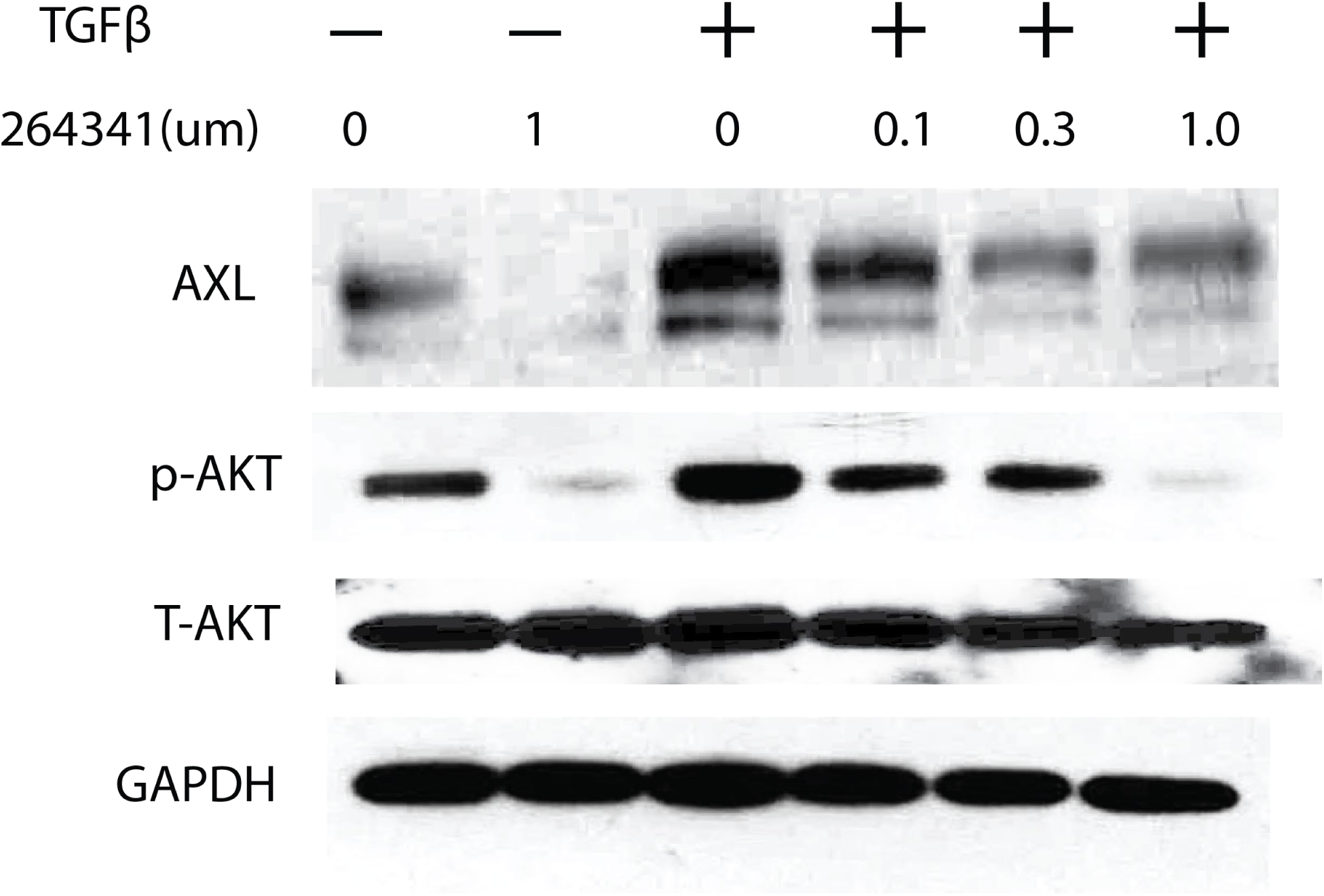
Western blot of CCD18co cell lysates treated with TGFβ and increasing doses of CCG264341. Protein expression of AXL and pAkt is shown, compared to total Akt and a GAPDH control in each lane. There is a dose-response reduction in AXL and pAkt.

**Figure 11.**
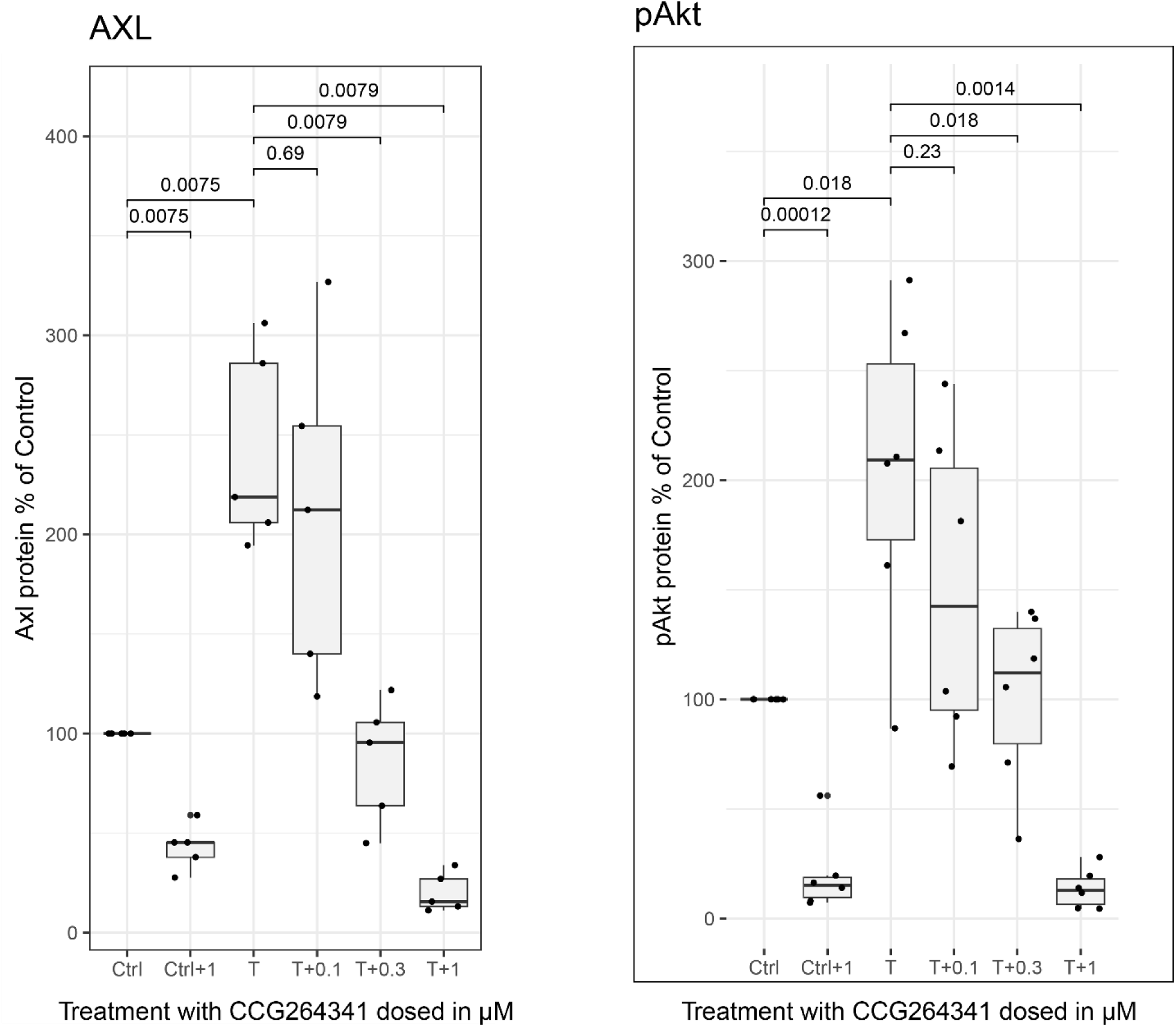
Plots of Western blot data for AXL and pAkt seen in Figure 10. CCD18co cell lysates treated with TGFβ and increasing doses of CCG264341. Protein expression of AXL and pAkt is shown, compared to total Akt and a GAPDH control in each lane. There is a dose-response reduction in AXL and pAkt.

**Figure 12.**
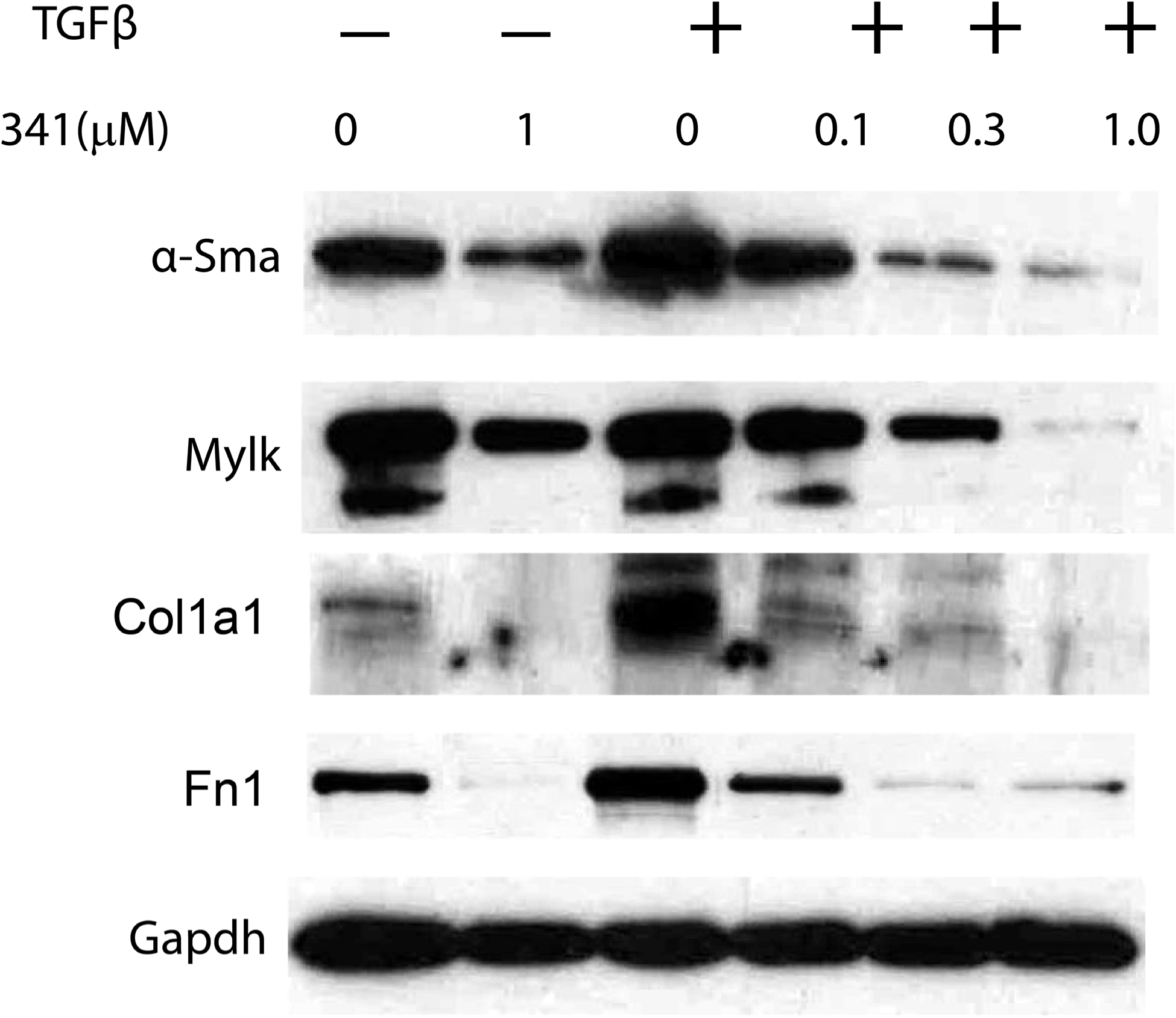
Western blot of αSMA, MYLK, Collagen 1, and fibronectin in CCD18co cell lysates treated with TGFβ and increasing doses of CCG264341. Higher doses of CCG264341 produce downstream effects similar to AXL knockdown with siRNA.

**Figure 13.**
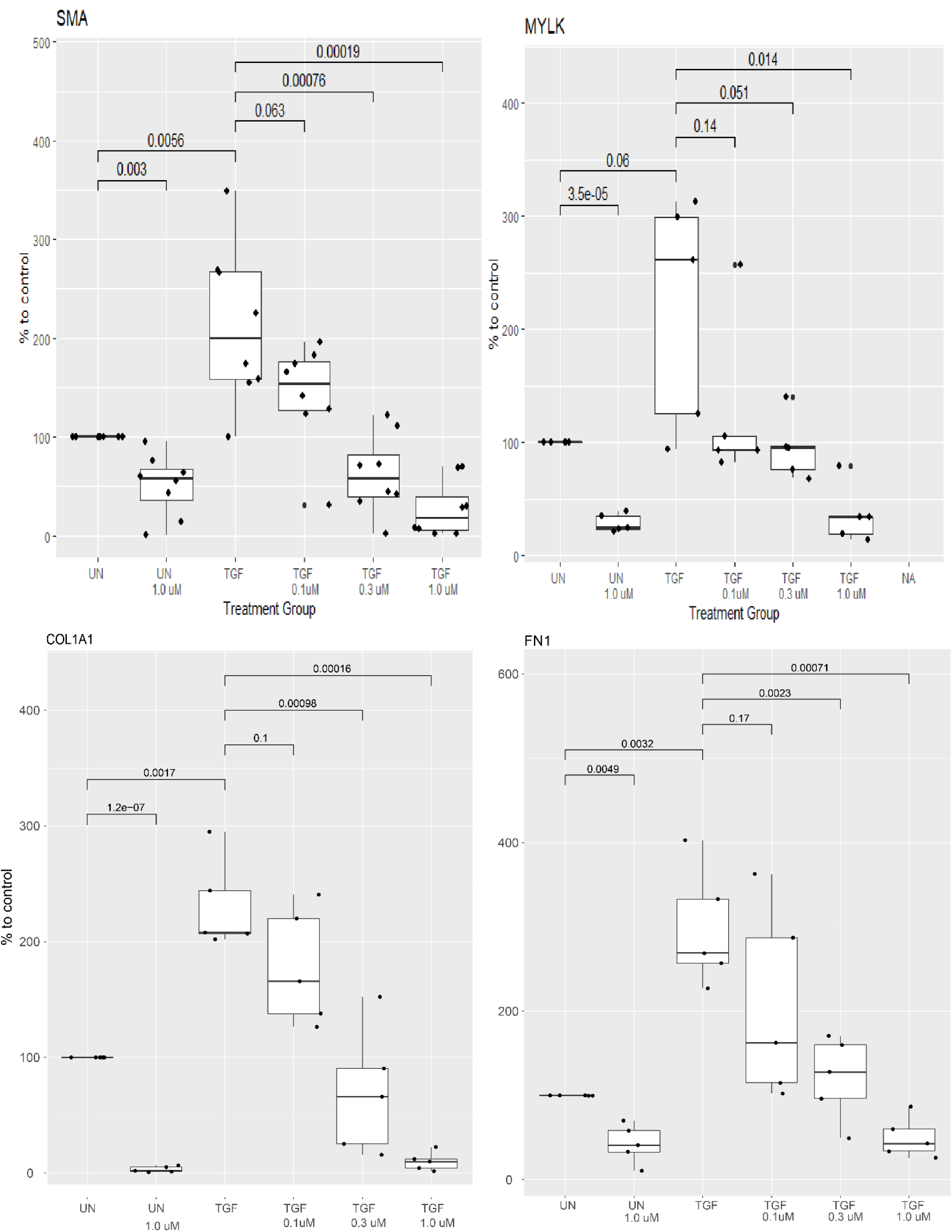
Plots of Western blot data for αSMA, MYLK, Collagen 1, and fibronectin seen in Figure 12.

**Figure 14.**
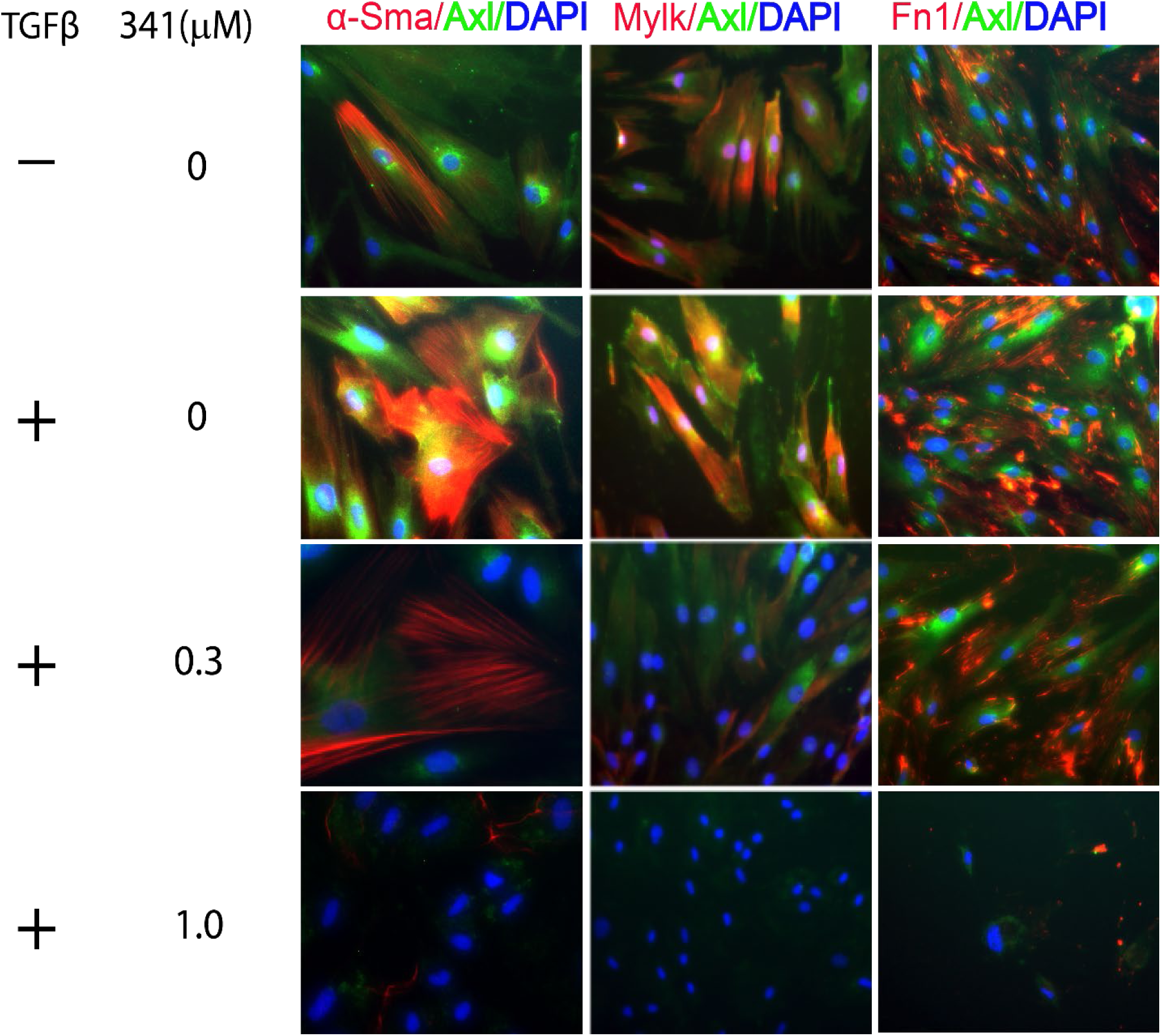
Immunofluorescence staining with αSMA, AXL, MYLK, DAPI, and fibronectin after treatment with TGFβ vs CCG264341 at varying doses

We then tested if Axl inhibition could lead to active apoptosis of myofibroblasts in the FasL cell model. These results showed that either knocking down AXL expression with siRNA (Figure 15) and inhibiting Axl activity with CCG264341 treatment could activate myofibroblast cell apoptosis as measured by increased cleaved-PARP expression, and that combining this further increased apoptosis.

**Figure 15.**
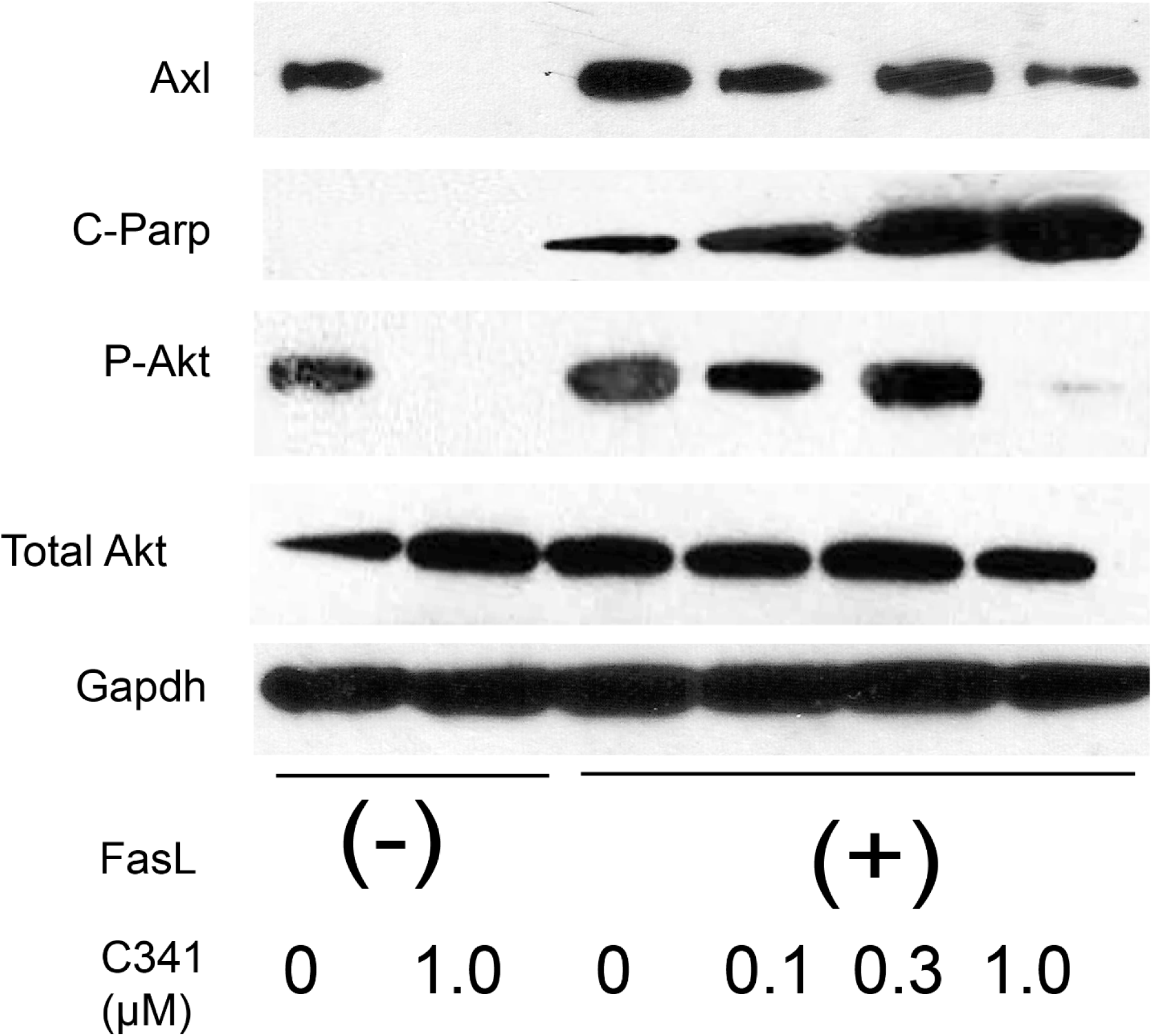
Apoptosis Effects in Human myofibroblasts with CCG264341 in the Fas ligand assay. CCG264341 appears synergistic in inducing apoptosis of CCD18co human myofibroblasts

These data demonstrate that CCG264341 does effectively block AXL activity *in vitro*, stimulates cell apoptosis, and suppresses fibrogenic pathways. The efficacy of CCG264341 *in vitro* was similar to AXL SiRNA knockdown, which supports the potency of this AXL inhibitor.

### CCG264341 treated intestinal fibrosis in the *Salmonella typhimurium* mouse model

We then wanted to evaluate whether CCG264341 could prevent fibrosis in vivo. First, we evaluated the concentration of CCG264341 in organs outside of the gut across a range of doses (Figure 16). Briefly,10/25/50/100 mg/kg of CCG264341 were given to mice via oral gavage (n=3-4 for each group) and organ tissue samples were collected 5 hours after treatment. The compound tissue concentrations showed that there is partial gut selectivity, especially in the terminal ileum, which could be especially valuable for human Crohn’s disease.

**Figure 16.**
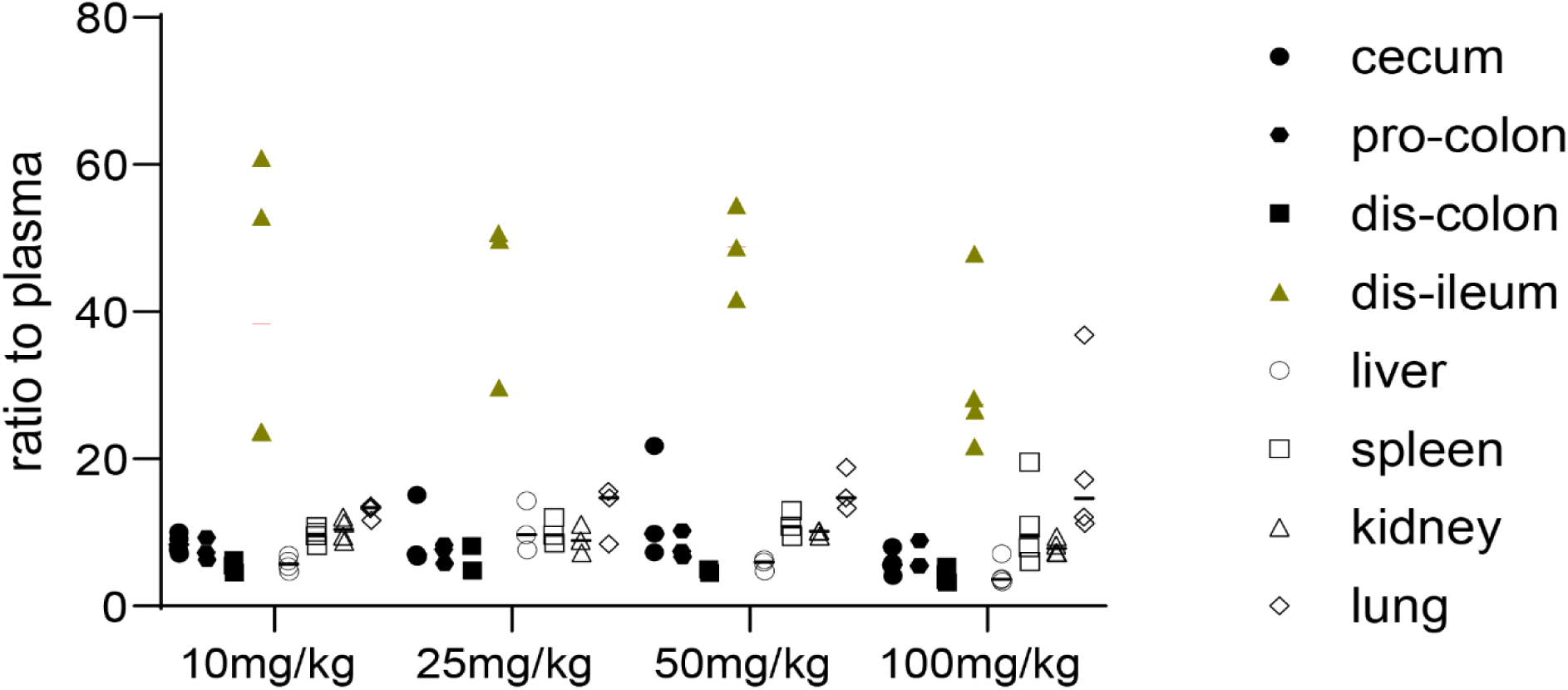
Tissue pharmacokinetics of CCG264341 across a range of doses and organs as measured in our PK Core.

As described in our previous work^25^*, Salmonella typhimurium* infected mice were used to test the anti-fibrotic effect of CCG264341, beginning treatment on day 5 after Salmonella infection and treating daily by oral gavage through day 21. Cecal and proximal colon fibrosis was observed in all of the *Salmonella typhimurium* infected mice by Masson’s trichrome staining. We used 2 doses of CCG264341, and 2.5 mg/kg/day treatment did not prevent fibrosis (data not shown), so we will focus on the 25 mg/kg/day CCG264341 treatment. Our data showed that *Salmonella* infection leads to mouse body weight loss (Figure 17, upper panel) and increased colon density due to a mixture of colon edema and fibrosis and shortening of the fibrotic colon (Figure 17, lower panel), while treatment with CCG264341 reduced body weight loss and colon density increases. CCG264341 concentrations were measured in control or *Salmonella typhimurium* infected mice after single or multiple doses (Figure 18), and these results showed partially gut selectivity, particularly in the terminal ileum, and that some drug accumulation in the gut tissues does occur with repeated dosing.

**Figure 17.**
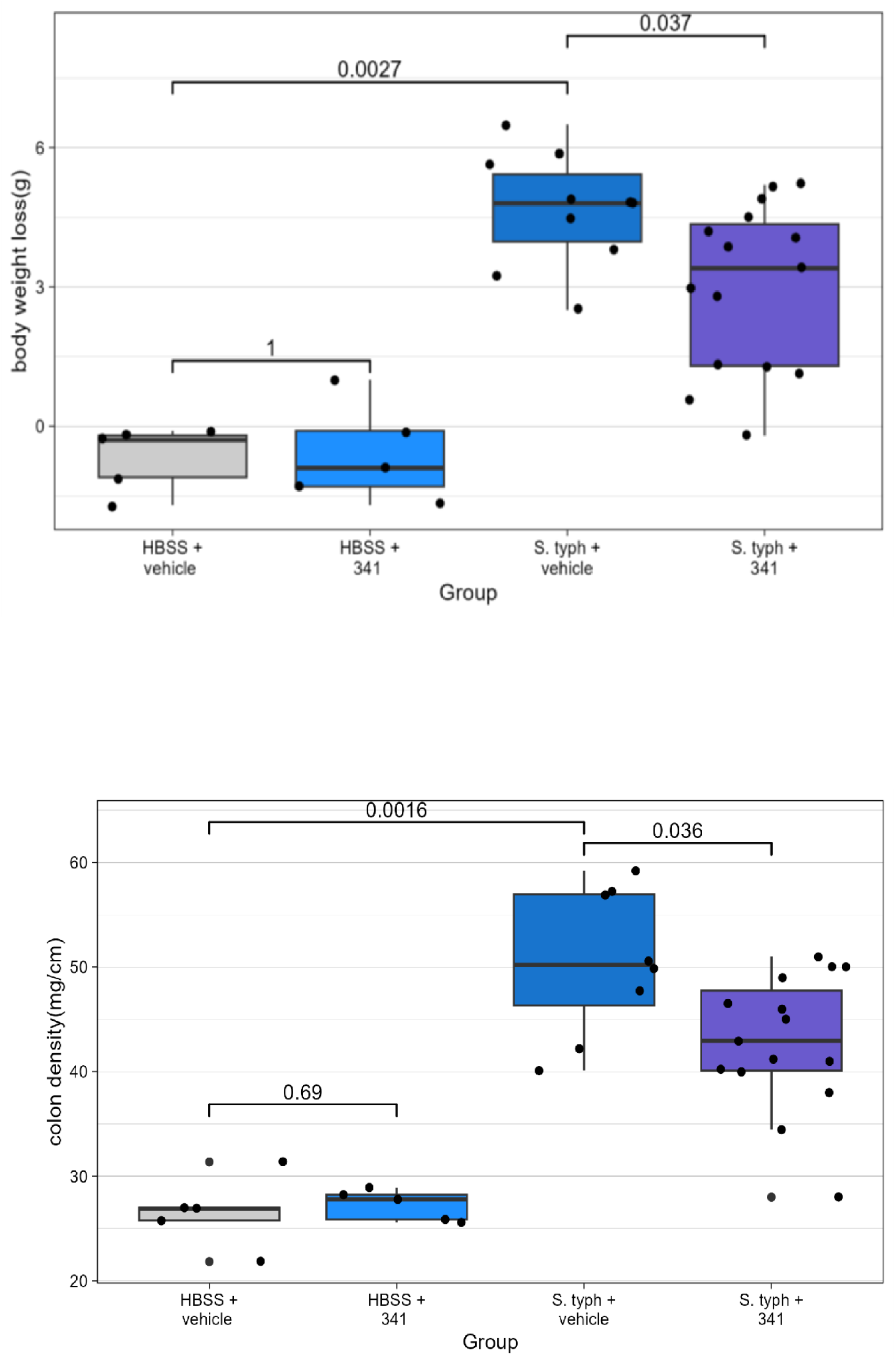
Effects of CCG264341 on Body Weight and Colon Density. Salmonella infection results in loss of mouse body weight and increased colon density as it becomes thicker and fibrotic and shorter, while CCG264341 partially reverses this effect.

**Figure 18.**
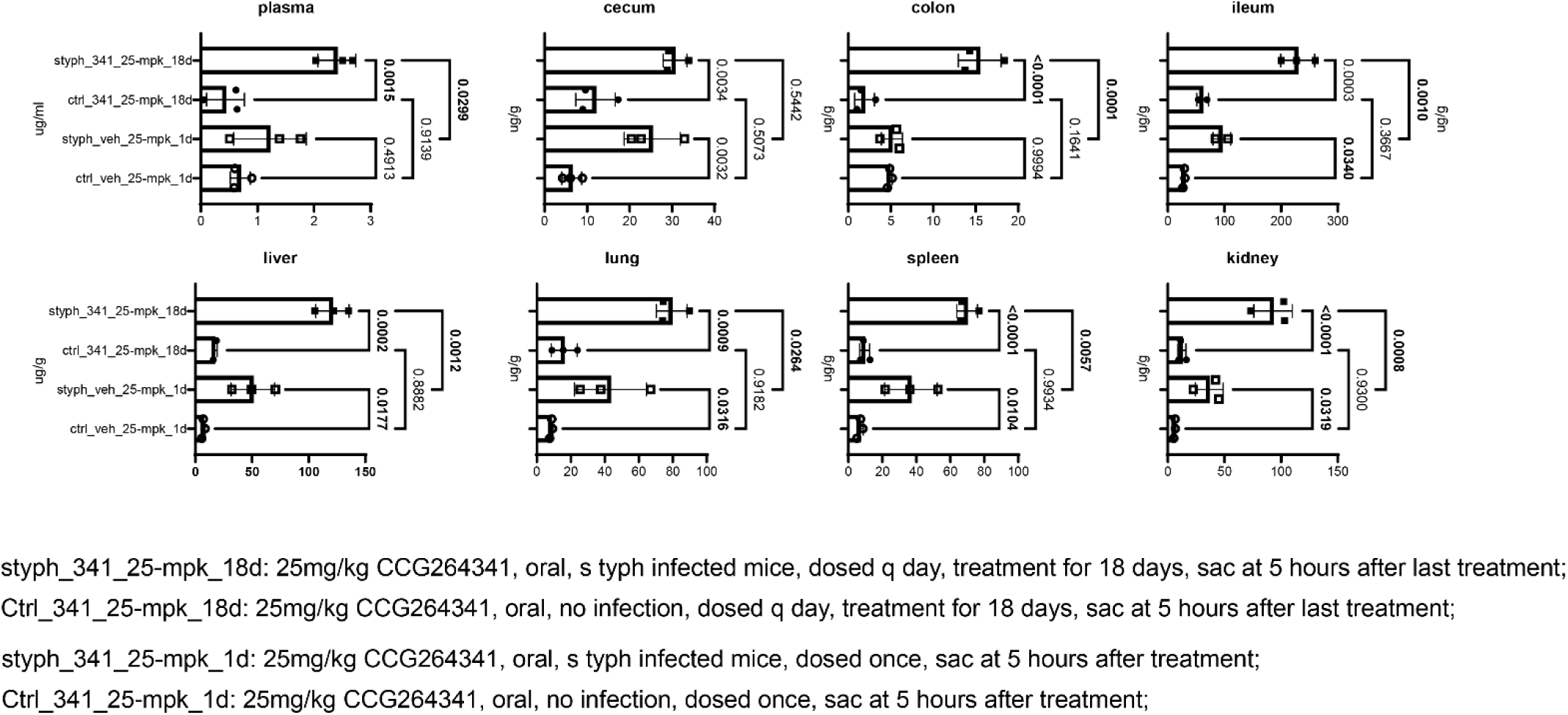
Single dose vs multi-day dosing of CCG264341 for PK. Gut selectivity and gut accumulation are seen over multiple daily doses.

AXL expression was increased in the colon with *Salmonella typhimurium* infection, and CCG264341 treatment reduced Axl expression to baseline levels (Figure 19). H&E staining results showed that *Salmonella typhimurium* infection caused significant increases in the blinded pathologist fibrosis score. In contrast, CCG264341 treatment of *Salmonella typhimurium* infected mice decreased the fibrosis score and significantly decreased the total histologic score (Figure 20 upper panels), Trichrome staining further confirmed this result. CCG264341 treatment decreased the fibrosis score and reduced the fibrotic area (Figure 20 lower panels).

**Figure 19.**
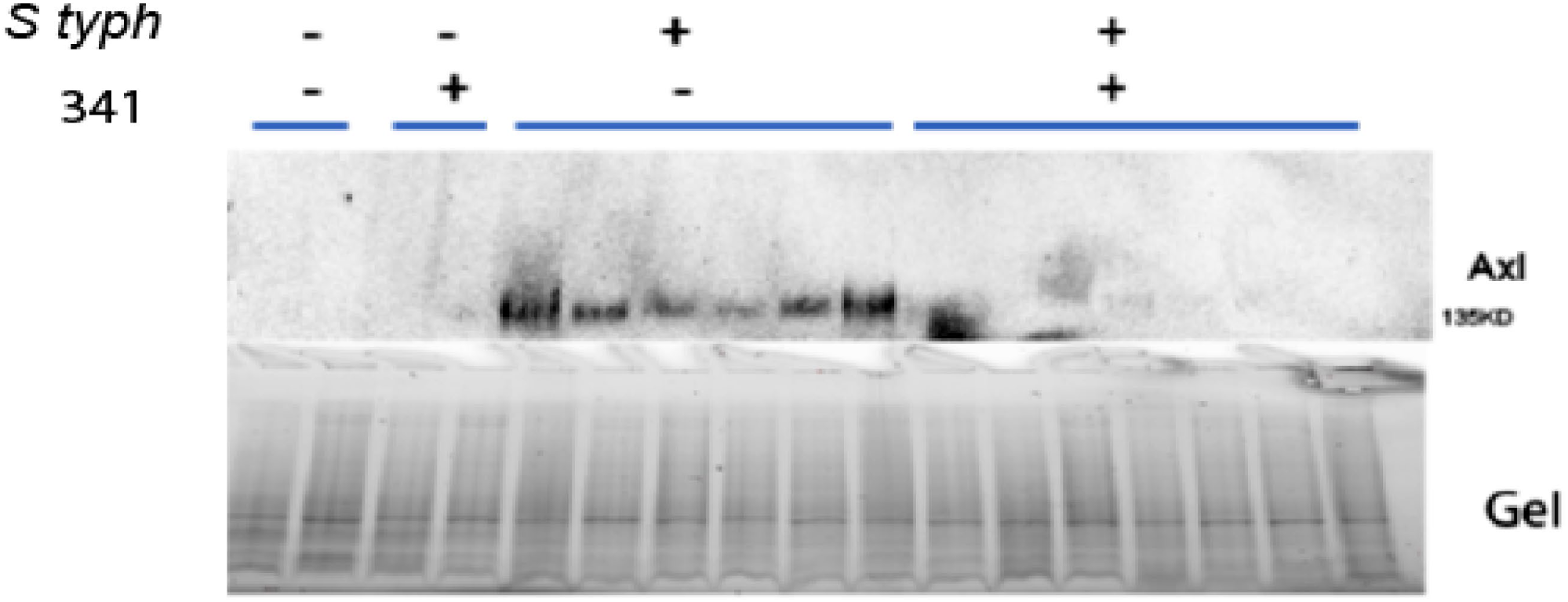
AXL protein is induced in mouse colon by Salmonella typhimurium infection and reduced by CCG264341. Western blot at top, protein gel for normalization below

**Figure 20.**
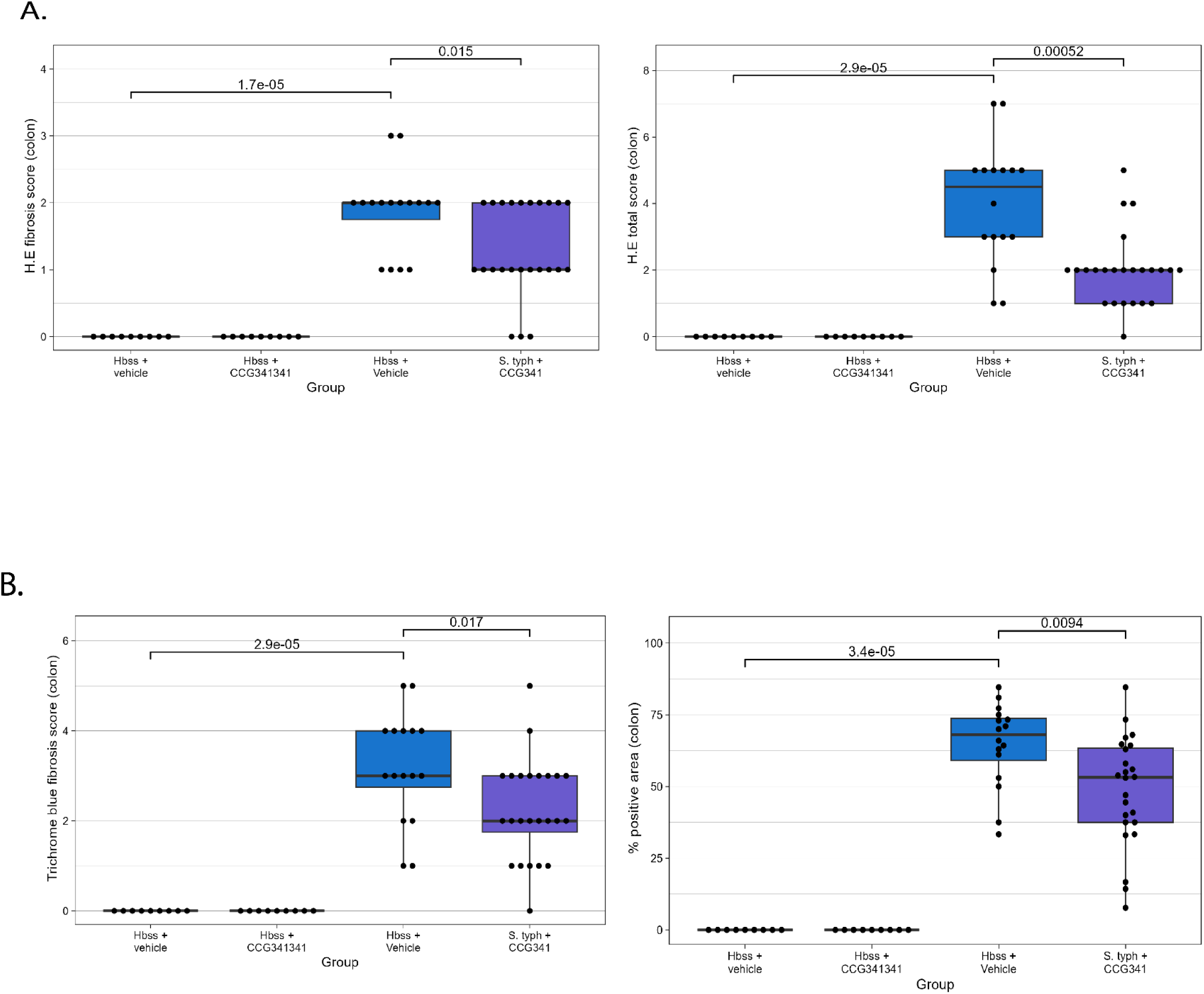
Blinded pathology scores for mouse colon for H&E Fibrosis, H&E total score, Trichrome fibrosis score, and Percent affected area for trichrome. CCG264341 treatment reduced fibrosis and the percentage area affected by fibrosis.

*Salmonella typhimurium* infection also increased Axl expression, and also elevated downstream signals p-Akt, p-Erk and p-STAT3 protein expression, and CCG264341 treatment reduced all of these signals (Figure 21). *Salmonella typhimurium* infection also caused an increased expression of fibrosis-related proteins, which were then reduced with CCG264341 treatment (Figure 21). immunofluorescence staining showed that CCG264341 treatment prevented the increases in protein expression of MYLK, FN1 and Col1a1 and decreased AXL expression in infected mice (Figure 22).

**Figure 21.**
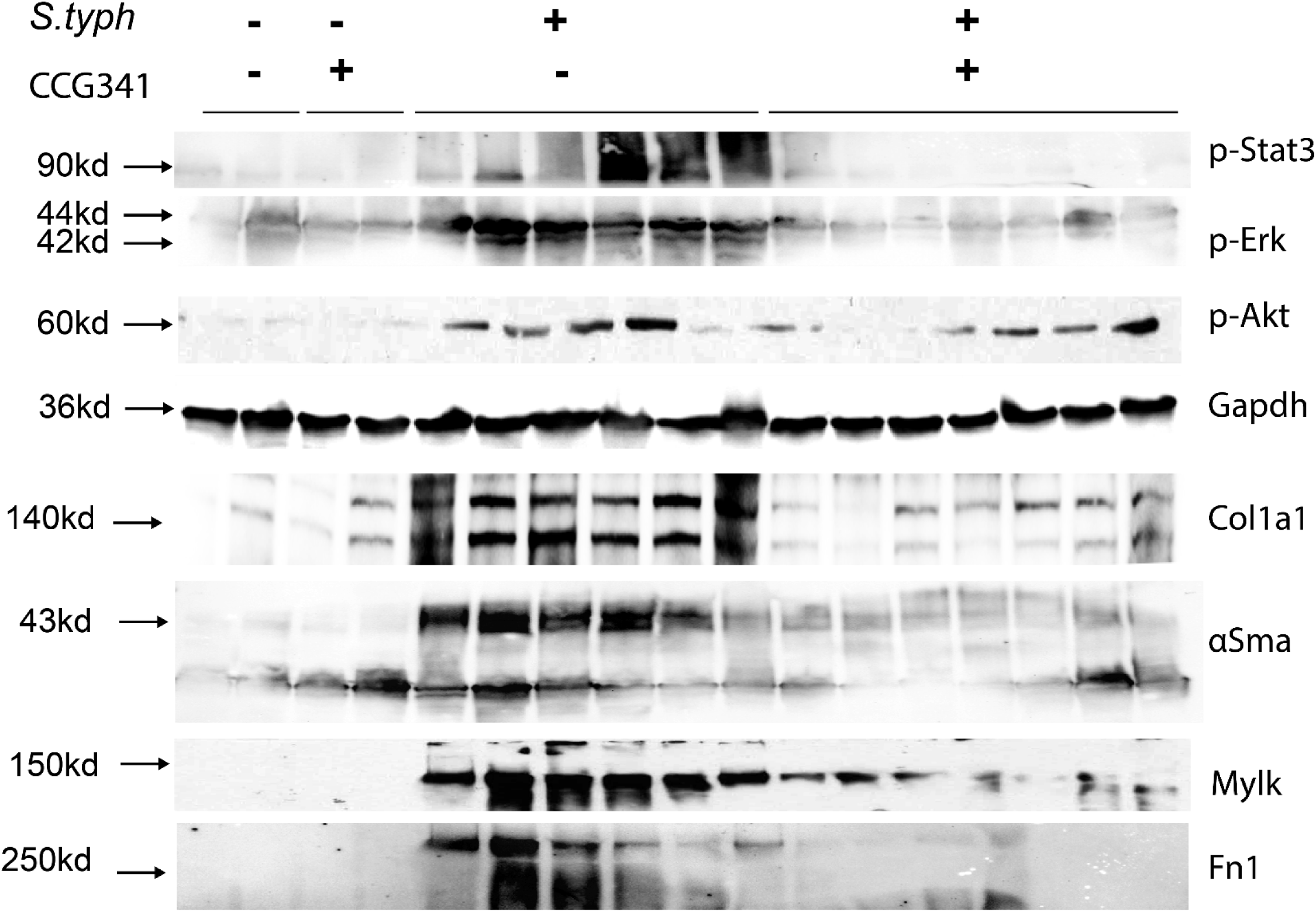
Western blot of mouse colon lysates without or with S. typhimurium infection and CCG264341 treatment at 25 mg/kg/day. CCG264341 reduces pSTAT3, pERK, pAkt, Collagen 1, aSMA, FN1 and MYLK after infection with *Salmonella typhi*.

**Figure 22.**
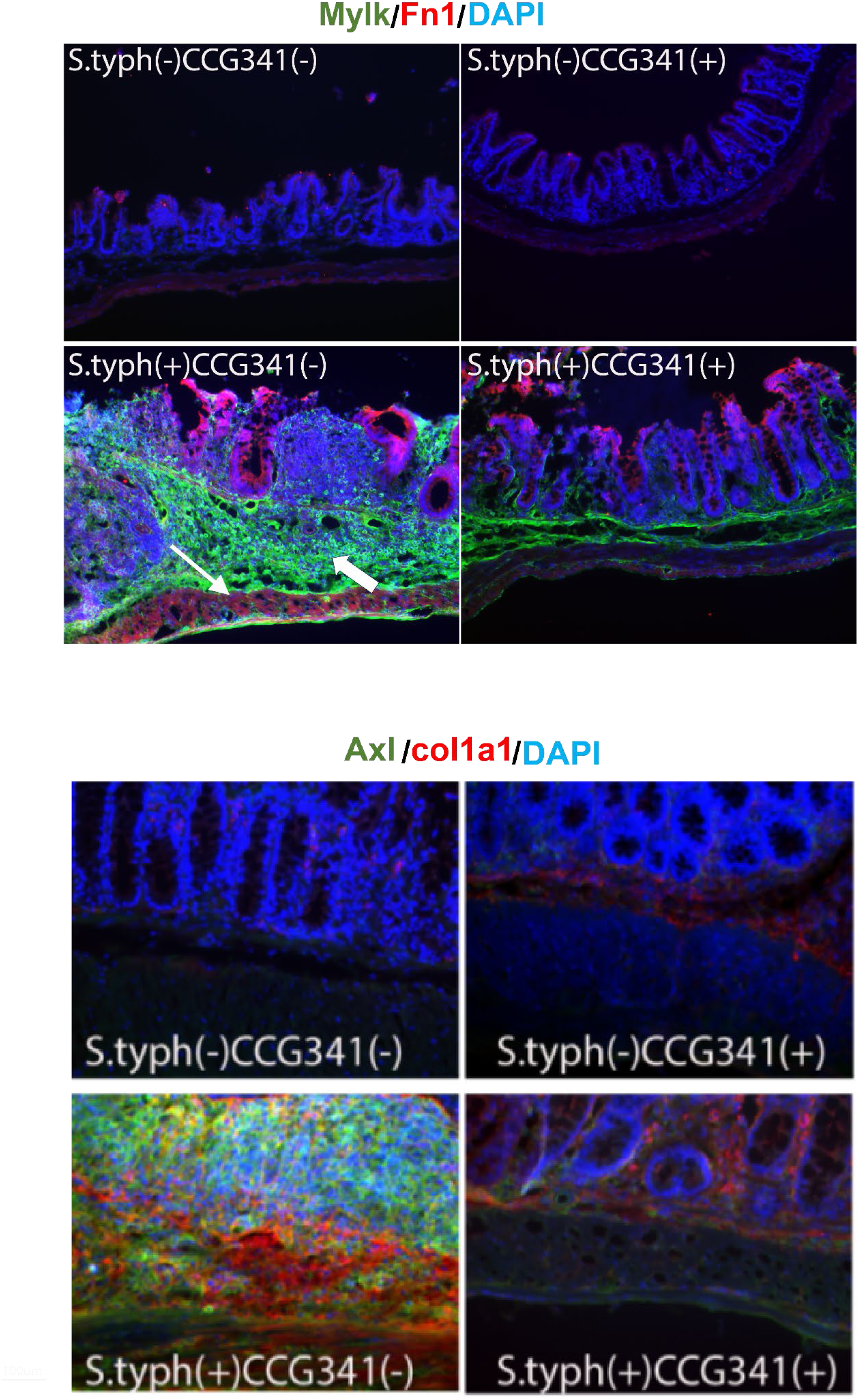
Immunofluorescence of mouse colon, stained with MYLK, Fibronectin, AXL, Collagen 1, and DAPI. Salmonella typhimurium significantly increases MYLK, fibronectin, AXL, and collagen 1, all of which are reduced by treatment with CCG264341.

Real-time PCR was performed to measure the colon cytokine expression (Figure 23), the results showed that *Salmonella typhimurium* infection caused increasing of TGFβ, IL1β, IL6 and INFϒ gene expression, CCG264341 treatment prevent these increasing, which may contribute to reduce the fibrosis.

**Figure 23.**
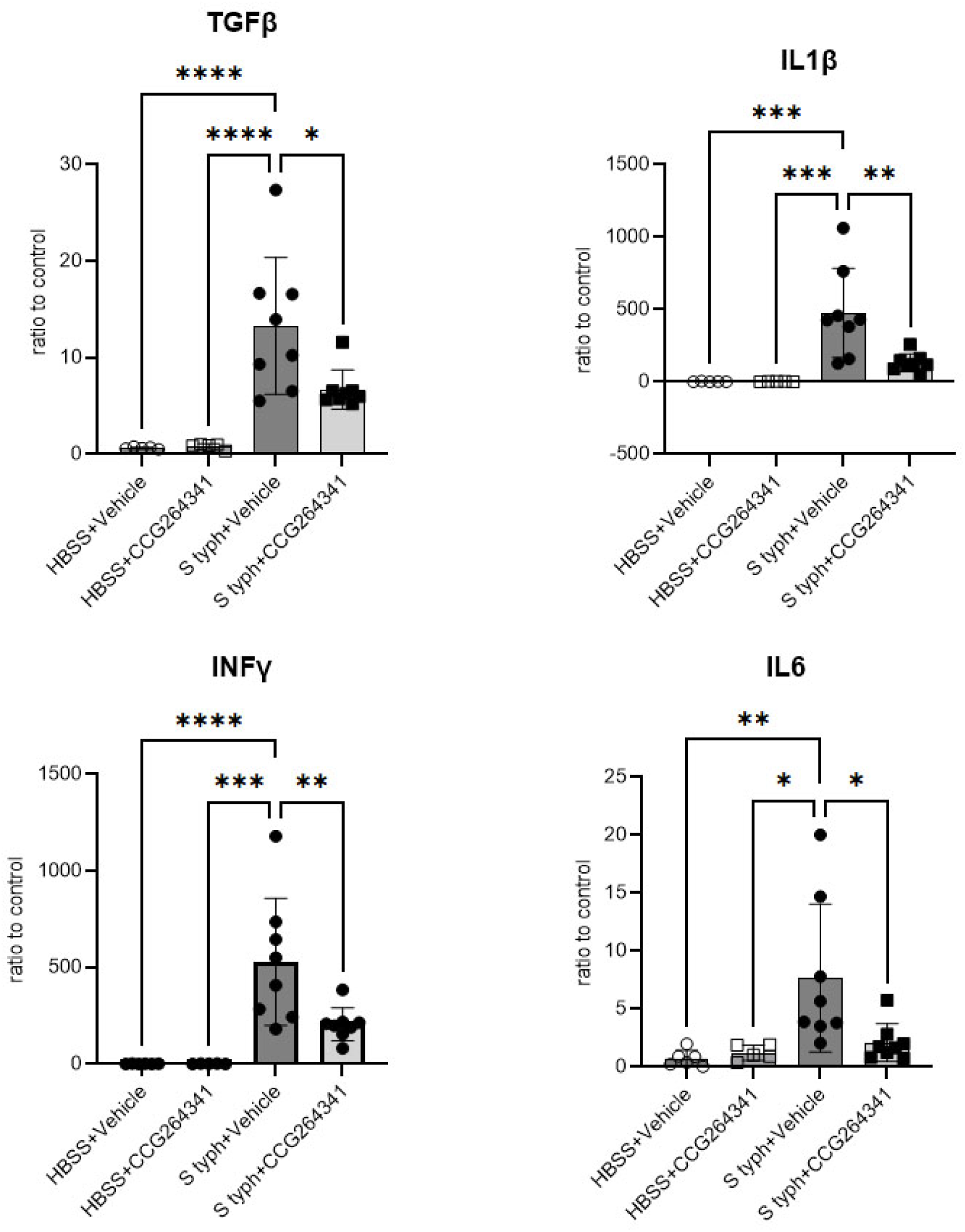
Inflammatory cytokine gene expression is also affected by CCG264341, with reduction from the high levels induced by infection with *Salmonella typhi*. Reductions in the gene expression of IL1β, IL6, TGFβ and IFNγ are evident.

## Discussion

A dreaded complication of Crohn’s disease is the development of obstructive fibrostenotic disease. There are currently no FDA approved medical therapies for the treatment of this complication, and surgery is ultimately required in as many as 64% of cases^27^ resulting in over 10 billion dollars in US healthcare costs annually^3,4^.

Our findings demonstrate that our novel Axl inhibitor CCG264341 with partial gut selectivity successfully treated intestinal fibrosis in a murine model with low toxicity. This is the first novel, gut selective AXL inhibitor to **effectively treat intestinal** fibrosis, and this study provides a new path toward effective and safe anti-fibrotic medical therapies for patients with structuring Crohn’s disease.

In this study, our PK data showed that very high compound concentrations could be achieved in the terminal ileum, which is the primary target organ in IBD patients with obstruction. There were also detectable levels of compound in the other high blood-flow organs like lung, kidney, and spleen compared to cecum and colon, which have led to off-target effects at 100 mg/kg dosing like splenomegaly (data not shown). Further improvement of this compound series to produce greater gut-selectivity will be a key to develop the next generation of candidate anti-fibrotic therapies.

As we know, Axl is expressed on the surface of different cells including high expression in macrophages^28–30^. In a limitation of this study, we focused on how this compound affects fibroblasts and fibrosis, but future studies on this candidate series will evaluate how this compound affects the immune system and inflammation, and the crosstalk between fibroblast and macrophage will be investigated.

In summary, fibrosis was observed in the colon and cecum of the mouse *S. typhimurium* model, with increased p-Akt/p-Erk/p-Stat3, and Mylk/Fn1/Col1a1/αSma, all downstream of AXL. Treatment with 25mg/kg/day oral CCG264341 partially prevented colon fibrosis by inhibiting AXL and blocking the activation of these downstream signaling pathways.

In conclusion, our results showed that CCG264341 was a partially gut selective AXL inhibitor. This compound effectively inhibited AXL expression i*n vitro* and partially reversed colon fibrosis in *Salmonella typhimurium* infected mice. Future work will aim to improve the gut-selectivity, which we hope will produce a stronger anti-fibrotic effect with minimal toxicity, with potential to treat the intestinal fibrosis of patients with Crohn’s disease.

